# Multi-Tissue Metabolomics Reveal mtDNA- and Diet-Specific Metabolite Profiles in a Mouse Model of Cardiometabolic Disease

**DOI:** 10.1101/2024.12.19.629398

**Authors:** Abhishek Shastry, Mia. S. Wilkinson, Dalia M. Miller, Michelle Kuriakose, Jennifer L.M.H. Veeneman, Matthew Ryan Smith, Charles C.T. Hindmarch, Kimberly J. Dunham-Snary

**Author notes:** Address for Correspondence: Kimberly Dunham-Snary, PhD Assistant Professor | Tier II Canada Research Chair Department of Biomedical & Molecular Sciences | Queen’s University Botterell Hall 429 | Kingston, ON | Canada | K7L 3N6.

## Abstract

**Rationale:** Excess consumption of sugar- and fat-rich foods has heightened the prevalence of cardiometabolic disease, which remains a driver of cardiovascular disease- and type II diabetes-related mortality globally. Skeletal muscle insulin resistance is an early feature of cardiometabolic disease and is a precursor to diabetes. Insulin resistance risk varies with self-reported race, whereby, African-Americans have a greater risk of diabetes development relative to their White counterparts. Self-reported race is strongly associated with mitochondrial DNA (mtDNA) haplogroups, and previous reports have noted marked differences in bioenergetic and metabolic parameters in cells belonging to distinct mtDNA haplogroups, but the mechanism of these associations remains unknown. Additionally, distinguishing nuclear DNA (nDNA) and mtDNA contributions to cardiometabolic disease remains challenging in humans. The Mitochondrial-Nuclear eXchange (MNX) mouse model enables *in vivo* preclinical investigation of the role of mtDNA in cardiometabolic disease development, and has been implemented in studies of insulin resistance, fatty liver disease, and obesity in previous reports.

**Methods:** Six-week-old male C57^nDNA^:C57^mtDNA^ and C3H^nDNA^:C3H^mtDNA^ wild-type mice, and C57^nDNA^:C3H^mtDNA^ and C3H^nDNA^:C57^mtDNA^ MNX mice, were fed sucrose-matched high-fat (45% kcal fat) or control diet (10% kcal fat) until 12 weeks of age (n = 5/group). Mice were weighed weekly and total body fat was collected at euthanasia. Gastrocnemius skeletal muscle and plasma metabolomes were characterized using untargeted dual-chromatography mass spectrometry; both hydrophilic interaction liquid chromatography (HILIC) and C18 columns were used, in positive- and negative-ion modes, respectively.

**Results:** Comparative analyses between nDNA-matched wild-type and MNX strains demonstrated significantly increased body fat percentage in mice possessing C57^mtDNA^ regardless of nDNA background. High-fat diet in mice possessing C57^mtDNA^ was associated with differential abundance of phosphatidylcholines, lysophosphatidylcholines, phosphatidylethanolamines, and glucose. Conversely, high-fat diet in mice possessing C3H^mtDNA^ was associated with differential abundance of phosphatidylcholines, cardiolipins, and alanine. Glycerophospholipid metabolism and beta-alanine signaling pathways were enriched in skeletal muscle and plasma, indicating mtDNA-directed priming of mitochondria towards oxidative stress and increased fatty acid oxidation in C57^nDNA^:C57^mtDNA^ wild-type and C3H^nDNA^:C57^mtDNA^ MNX mice, relative to their nDNA-matched counterparts. In mtDNA-matched mice, C57^mtDNA^ was associated with metabolite co-expression related to the pentose phosphate pathway and sugar-related metabolism; C3H^mtDNA^ was associated with branched chain amino acid metabolite co-expression.

**Conclusions:** These results reveal novel nDNA-mtDNA interactions that drive significant changes in metabolite levels. Alterations to key metabolites involved in mitochondrial bioenergetic dysfunction and electron transport chain activity are implicated in elevated beta-oxidation during high-fat diet feeding; abnormally elevated rates of beta-oxidation may be a key driver of insulin resistance. The results reported here support the hypothesis that mtDNA influences cardiometabolic disease-susceptibility by modulating mitochondrial function and metabolic pathways.

## Introduction

The prevalence of cardiometabolic disease (CMD) has increased globally over the last three decades ^1–3^. CMD is an array of metabolic abnormalities characterized by the presence of three or more of the following factors: abdominal obesity, high blood triglycerides, low HDL cholesterol, high blood pressure, and high fasting blood glucose ^4^. If left unmanaged, CMD increases the risk for type 2 diabetes mellitus (T2DM) and cardiovascular disease, among the leading causes of death worldwide ^2,5–8^. While several strategies to mitigate CMD risk factors are available, including dietary, exercise, and pharmaceutical interventions ^9^, none of these have reversed the global trajectory of CMD. According to World Obesity Atlas projections, 1 in 4 adults and, alarmingly, 1 in 5 children and adolescents will be obese by 2035. Furthermore, global expenditure related to obesity is set to increase from $15.2 billion in 2020 to $23.0 billion in 2035^10^.

In addition to environmental and lifestyle factors, the underlying genetic drivers of disease remain elusive in large populations. In North America, African-Americans, African-Canadians, and North American Indigenous populations are at greater risk for T2DM compared to their White (Western Europe-descended) counterparts ^11,12^. Interestingly, these disparities persist after controlling for CMD-associated behaviours (e.g., smoking, alcohol consumption), socioeconomic status, and medication status ^11,13–19^. Genome-wide association studies (GWAS) of nuclear DNA (nDNA) have uncovered >1000 single-nucleotide polymorphisms (SNPs) associated with obesity and other CMD abnormalities ^20,21^. Several of these SNPs, including proprotein convertase subtilisin/kexin type 9 (PCSK9), fat mass and obesity-associated protein (FTO), and tub bipartite transcription factor (TUB), have been identified in African-Americans and may contribute to increased CMD rates ^22–24^. However, significant heterogeneity exists within self-reported racial groups and subgroups which affects SNP validity and reproducibility ^25^.

While the mammalian nuclear genome encodes ∼20,000 genes ^26^, eukaryotic cells also contain DNA encoding a second genome situated in the mitochondria. Mitochondrial DNA (mtDNA) encodes 2 rRNAs, 22 tRNAs, and 13 mRNAs, which are used to generate the structural and catalytic polypeptide subunits of electron transport chain complexes ^27^. Phylogenomic analyses demonstrate that the mutational rate of nDNA is significantly lower than that of mtDNA ^28,29^. mtDNA mutations can clonally expand in downstream generations, increasing the mutational burden of pathological mtDNA variants ^30^. As such, SNPs in the mitochondrial genome have been investigated for their contribution to various diseases ^31^. Point mutations in mtDNA genes encoding tRNA^Leu^, tRNA^Lys^, tRNA^Ser^, and ATP6 have been associated with diabetes phenotypes^32^. As a protective mechanism, bottleneck effects during oocyte development prevent the germline transmission and increased heteroplasmy of pathological mtDNA transmission to offspring ^33^; however, mutations that appear to be immediately non-pathological and do not affect the fitness or survival of the organism may still be inherited. mtDNA variants common to humans whose ancestors originated in geographically isolated regions are clustered into haplogroups ^34–37^. It has been demonstrated that humans belonging to distinct mtDNA haplogroups display varied levels of oxidative stress and ATP production efficiency ^38^. However, the mtDNA- and nDNA-specific contributions to these bioenergetic profiles have largely been uncharacterized.

Skeletal muscle is a highly metabolically active tissue whose function is greatly influenced by close systemic regulation of insulin signalling ^39,40^. Skeletal muscles are composed of fibers that can be broadly grouped into two types: Type I fibers (“slow twitch”) are mitochondrially dense and produce a large amount of energy over a long term through oxidative metabolism; and Type II fibers (“fast twitch”), which largely rely on glycolytic metabolism for short bursts of energy through rapid ATP production ^39^. As such, disruptions in glucose import into skeletal muscle can cause dysfunction in muscle primarily containing fast twitch fibers ^41^. The excessive consumption of diets high in fat is known to drive insulin resistance, especially in insulin-sensitive tissues ^42–44^. Prolonged high-fat diet (HFD) consumption is known to cause mitochondrial dysfunction, both morphologically (i.e., abnormal fusion and fission) and bioenergetically through dysregulation of complete fatty acid oxidation ^45,46^. Furthermore, evidence from multiple sources indicates that mitochondrial genetic background can modulate insulin sensitivity systemically and in skeletal muscle tissue ^47–50^. Additionally, HFD and pathological mtDNA variants influence global gene and metabolite expression profiles in metabolically active tissue types such as white adipose and skeletal muscle ^51–54^. However, the influence of a HFD on the underlying skeletal muscle metabolites of organisms belonging to distinct mitochondrial genetic backgrounds is unclear. Fluctuations in critical metabolites involved in mitochondrial bioenergetics and metabolism in the context of CMD have been reviewed previously ^55^.

Disentangling the impact of mtDNA from nDNA is inherently challenging due to their interdependent functions; mtDNA encodes a bacterial genome that has evolved as an endosymbiont ^56,57^, and its 37 genes interact with over 20,000 nuclear genes. This complexity is heightened when studying mtDNA effects in adult humans, as mtDNA lacks the protective histones and comprehensive repair mechanisms of nDNA ^58–63^. The ability to effectively ‘silence’ nDNA effects is largely achieved in the Mitochondrial-Nuclear eXchange (MNX) mouse model. MNX mice are generated through reciprocal pronuclear exchange between two wild-type (WT) inbred mouse strains in embryo ^64^; this process results in mice that possess native nDNA but mismatched mtDNA backgrounds compared to their WT counterparts. Through pairwise comparisons between nDNA-matched WT and MNX mice, this model allows for the investigation of mtDNA background-dependent modulation of disease phenotypes.

In studies of CMD, C57BL/6J and C3H/HeN inbred mice are used in the MNX model due to their known differing susceptibility to CMD-associated phenotypes, including systemic insulin resistance, hepatic inflammation, reactive oxygen species (ROS) production, and atherosclerosis ^65–72^. Previous research in MNX mice has demonstrated exacerbated liver fibrosis and inflammation, pathological adipose and hepatic gene expression, reduced circulating and skeletal muscle glucose tolerance, and greater oxidative stress in cardiac overload in mice harboring C57^mtDNA^ relative to C3H^mtDNA^, independent of nDNA background ^47,73–75^. Other MNX models using FVB and BALB/c mice, have demonstrated tumour growth and localization patterns that are modulated by mtDNA background ^76,77^.

Previous studies of MNX mice reveal that mice possessing C57^mtDNA^ exhibit pathological phenotypes in numerous tissues, including the skeletal muscle ^78^. The metabolomic landscape of early CMD in mice with distinct mtDNA backgrounds is currently undescribed, and changes in metabolite levels may influence CMD susceptibility in response to dietary stress. The present study evaluated skeletal muscle and plasma metabolites that are altered in HFD-induced CMD using the MNX mouse model. We hypothesize that mice harboring C57^mtDNA^ would exhibit metabolite levels reflective of lipid-induced insulin resistance in skeletal muscle and metabolites reflective of impaired multi-organ communication in plasma, relative to mice harboring C3H^mtDNA^.

## Methods

### Animal Ethics

All animal studies were performed in accordance with protocols approved by the University Animal Care Committee of Queen’s University at Kingston (Protocols #2021-2156 and #2023-2390).

### Mice

Wild-type C57BL/6J (C57^nDNA^:C57^mtDNA^) mice were purchased at 3-4 weeks of age (Jackson Laboratories, Bar Harbor, ME). Wild-type C3H/HeN (C3H^nDNA^:C3H^mtDNA^) and MNX (C57^nDNA^:C3H^mtDNA^ and C3H^nDNA^:C57^mtDNA^) mice were generated from an in-house colony maintained at Queen’s University. Briefly, female MNX mice are crossed with WT male mice of matching nDNA (e.g., a C57^nDNA^:C57^mtDNA^ WT male sires with C57^nDNA^:C3H^mtDNA^ MNX females); due to the sole maternal inheritance of mtDNA, all progeny of these breeding pairs are therefore MNX mice. The methods for the original generation of MNX mouse strains have been previously described ^52,64,75^. An overview of the MNX mouse model is presented in **Supplementary Figure S1**.

### Haplotyping

Using C57BL/6J mice as a reference genome, C3H/HeN mice exhibit non-synonymous mutations at nucleotide 9461 (T/A → C) encoding NADH dehydrogenase subunit 3 (Nd3) and nucleotide 9348 (G → A) encoding cytochrome c oxidase III (Cox3); additionally, C3H/HeN mice possess a ‘TA’ insert at nucleotide 9818, within tRNA^Arg^ ^79^. Wild-type (WT) and MNX mice had mitochondrial haplotypes confirmed using restriction fragment length polymorphism analysis based on known mtDNA SNPs, as described ^52,64^ (**Supplementary Figure S2**).

### Animal Diets

3-4-week-old male C57^nDNA^:C57^mtDNA^ WT, C3H^nDNA^:C3H^mtDNA^ WT, C3H^nDNA^:C57^mtDNA^ MNX, and C57^nDNA^:C3H^mtDNA^ MNX mice (n = 5/strain/diet) were weaned onto standard chow diet (LabDiet^®^ 5001, LabDiet, St. Louis, MO). At 6 weeks of age, mice were randomized into groups and provided either control diet containing 10% kcal fat (D12450H, Research Diets, New Brunswick, NJ) or sucrose-matched high-fat diet (HFD) containing 45% kcal fat (D12451, Research Diets) *ad libitum* for six weeks.

### Euthanasia

Mice were anesthetized using 5% inhaled isoflurane, transferred to surgical pads and maintained at the surgical plane of anesthesia using 3-4% isoflurane. When unresponsive to stimuli, mice were euthanized via cardiac puncture, and whole blood was collected and preserved using 500 units/mL heparin sodium salt (H3149-50KU, Sigma-Aldrich) in PBS (10010049, Gibco) to prevent clotting.

### Body Weight and Fat Pad Mass

Animal weight was monitored weekly from commencing diet exposure until euthanasia. All adipose tissue depots were dissected at euthanasia and weighed. Percentage body fat measurements are expressed as mass of total adipose tissue per total body weight.

### Plasma and Skeletal Muscle Collection

Whole blood was collected in syringes filled with 0.1 mL heparin as described above. Blood was centrifuged at 2000 × *g* for 15 min at room temperature to isolate plasma. Plasma was flash-frozen in liquid nitrogen and volumes were recorded. Gastrocnemius skeletal muscle, which is largely composed of type IIB fibres, was removed from hindlimb ^80,81^. Muscles were collected, weighed, and subsequently flash-frozen in liquid nitrogen.

### High Performance Liquid Chromatography-Mass Spectrometry (HPLC-MS)

Plasma and skeletal muscle samples were shipped to the Clinical Biomarkers Laboratory at Emory University (Atlanta, GA, USA) on dry ice. Samples were randomized to balance strain and diet between batches. Samples were preprocessed, analyzed using a dual-column HPLC-MS method (HILIC under positive electrospray ionization and reverse-phase C18 under negative electrospray ionization), and mass spectral data was extracted as described ^82–85^. For downstream analyses, peak intensities were quantile-normalized and log-transformed separately for skeletal muscle and plasma datasets. Features tables were filtered two-fold: (1) Features with zero counts in more than 20% of the samples across each dataset were removed; and (2) features with zero counts in more than 50% of the samples within each strain-diet group were removed. Using xMSannotator ^86^, only metabolites with identity confidence scores > 0 and primarily containing an “M+H” or “M-H” adduct were matched to features; annotations were made using the Human Metabolome Database (HMDB) ^87^ at a tolerance of 5 ppm. Unknown features were annotated manually through consultation with HMDB.

### Partial Least Squares-Discriminant Analysis (PLS-DA)

Sparse partial least squares-discriminant analysis (sPLS-DA) was performed in nDNA-matched mice in skeletal muscle and plasma, using the R package mixOmics v6.28.0, with 100 × 5-fold cross-validation ^88^. sPLS-DA is a supervised learning method that uses data with class labels to maximize separation between classified groups. The sparse algorithm was tuned after identifying the optimal number of principal components to minimize false classification while encapsulating the largest co-variance differences between groups ^89^.

### Differential Abundance Analysis

Pairwise contrasts were performed using Limma v3.60.2 ^90^ between control and HFD groups within each nDNA-matched strain in skeletal muscle and plasma. *P*-values were adjusted using the Benjamini-Hochberg method for multiple tests (*P*-adj) ^91^. *P*-adj < 0.2 was considered statistically significant, which is considered stringent in previous works ^83,84,92,93^.

### Weighted Metabolite Co-Expression Network Analysis

To evaluate metabolite co-expression patterns associated with the response to HFD in nDNA-match strains, a novel two-dimensional Weighted Gene Co-Expression Network Analysis (WGCNA) package called multiWGCNA was applied ^94^. Quantile-normalized, log-transformed peak intensities were used to adapt multiWGCNA for metabolomics. To reduce the dimensionality of the final matrix, the top 2,500 metabolites were selected by variance across skeletal muscle and plasma datasets separately. Skeletal muscle and plasma networks were constructed using the following parameters: minimum module size = 40, minimum KME to stay = 0.7, merge cut height = 0, power = 10, maxBlockSize = 25000).

multiWGCNA allows for the construction of networks across two experimental factors; diet was the primary factor, and strain was the secondary factor. As HFD feeding is predicted to be associated with CMD development, we investigated metabolites within HFD networks that were differentially co-expressed in WT and MNX strains ^95,96^. Overlap analysis was performed to evaluate HFD modules that were significantly associated (*P*-adj < 0.05) with strain modules with > 1 metabolite overlap. Subsequently, module preservation analysis incorporating a permutation test (1000 permutations) was performed to identify HFD modules not preserved in the control diet network (after supplying randomized class labels). The *Z*-summary score is a summary of module preservation, such that *Z*-summary < 10 indicates a lower probability of preservation and that the HFD module is associated specifically with HFD-induced CMD ^94^. HFD modules associated with WT and MNX strains were then filtered by the list of unpreserved HFD modules. As metabolite co-expression in response to HFD may induce similar pathological responses in WT and MNX mice, only HFD modules associated with WT or MNX strains (not both) were retained. This filtered list of HFD modules was annotated using HMDB as mentioned above.

### Pathway Enrichment Analysis

Pathway Enrichment Analysis was performed for all analyses using the Kyoto Encyclopedia of Genes and Genomes (KEGG) *Mus musculus* library in MetaboAnalyst 5.0 ^97,98^. Pathways were considered significant if *P* < 0.05.

### Other Statistical Analyses

Data for weekly body weight measurements and total body fat percentage are expressed as mean ± SEM. The significance of differences between baseline and six weeks of control and HFD in all strains was analyzed using two-way ANOVA, with Tukey post hoc analysis applied for multiple group comparisons using GraphPad Prism v10.2.3 (GraphPad, La Jolla, CA). Tests were considered significant when *P* < 0.05.

## Results

### Total Body Fat Segregates with mtDNA Signature

Total body fat percentage is a better predictor of early insulin resistance compared to body weight indices ^99^; mtDNA modulates total body fat percentage and weight change after six weeks of diet exposure, concordant with previous reports ^52^ **(Figure 1).**

**Figure 1:**
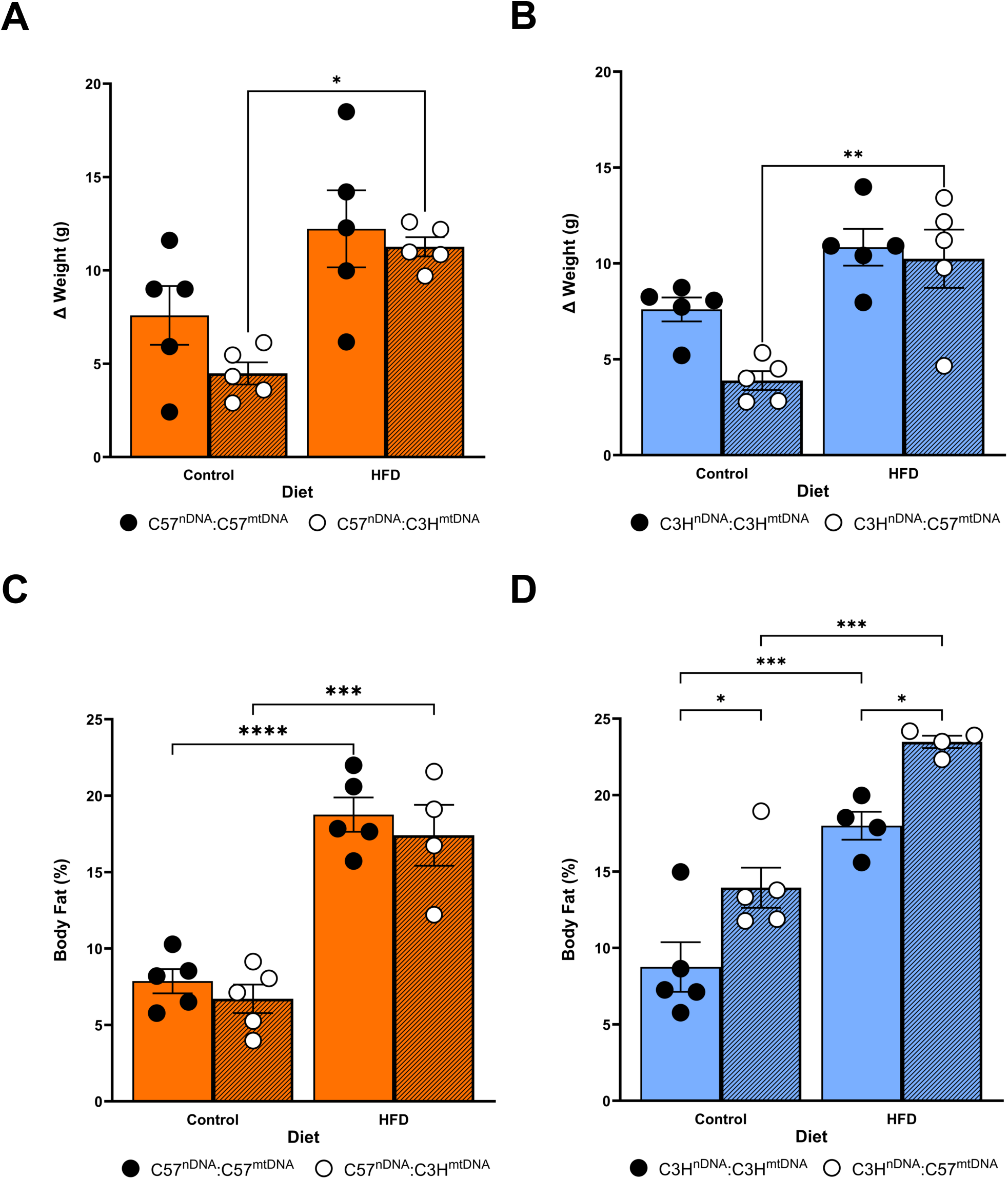
Morphometric measures of weight gain and fat mass in nDNA-matched mice fed Control or High-Fat Diet (HFD) for 6 weeks. **(A)** Difference in weight from 0 weeks to 6 weeks on diet in Control Diet-fed C57^nDNA^:C57^mtDNA^ WT (closed circle) and C57^nDNA^:C3H^mtDNA^ MNX (open circle) and HFD-fed C57^nDNA^:C57^mtDNA^ WT (closed circle) and C57^nDNA^:C3H^mtDNA^ MNX mice (open circle). **(B)** Difference in weight from 0 weeks to 6 weeks on diet in Control Diet-fed C3H^nDNA^:C3H^mtDNA^ WT (closed circle) and C3H^nDNA^:C57^mtDNA^ MNX (open circle) and HFD-fed C3H^nDNA^:C3H^mtDNA^ WT (closed circle) and C3H^nDNA^:C57^mtDNA^ MNX mice (open circle). **(C)** Total body fat percentage at sacrifice (6 weeks on diet) in Control Diet-fed C57^nDNA^:C57^mtDNA^ WT (closed circle) and C57^nDNA^:C3H^mtDNA^ MNX (open circle) and HFD-fed C57^nDNA^:C57^mtDNA^ WT (closed circle) and C57^nDNA^:C3H^mtDNA^ MNX mice (open circle). **(D)** Total body fat percentage at sacrifice (6 weeks on diet) in Control Diet-fed C3H^nDNA^:C3H^mtDNA^ WT (closed circle) and C3H^nDNA^:C57^mtDNA^ MNX (open circle) and HFD-fed C3H^nDNA^:C3H^mtDNA^ WT (closed circle) and C3H^nDNA^:C57^mtDNA^ MNX mice (open circle). Asterisks * indicate a significant difference exists between indicated groups, \**P* < 0.05, \*\*\**P* < 0.001, \*\*\*\**P* < 0.0001, ns – not significant, (n = 4-5/group).

### sPLS-DA Reveals Strain- and Diet-Dependent Principal Components

#### Skeletal Muscle

Control diet groups in C57^nDNA^:C57^mtDNA^ WT and C57^nDNA^:C3H^mtDNA^ MNX skeletal muscle showed clear separation along the first principal component (PComp), whereas HFD groups showed large separation in the third PComp. The third PComp also displayed the greatest separation between HFD and control groups (**Figure 2A**). There was no separation of control diet groups in C3H^nDNA^:C3H^mtDNA^ WT and C3H^nDNA^:C57^mtDNA^ MNX skeletal muscle; however, HFD groups separated along the third PComp. Similar to mice harboring C57^nDNA^, HFD and control groups separated along the third PComp (**Figure 2B**).

**Figure 2:**
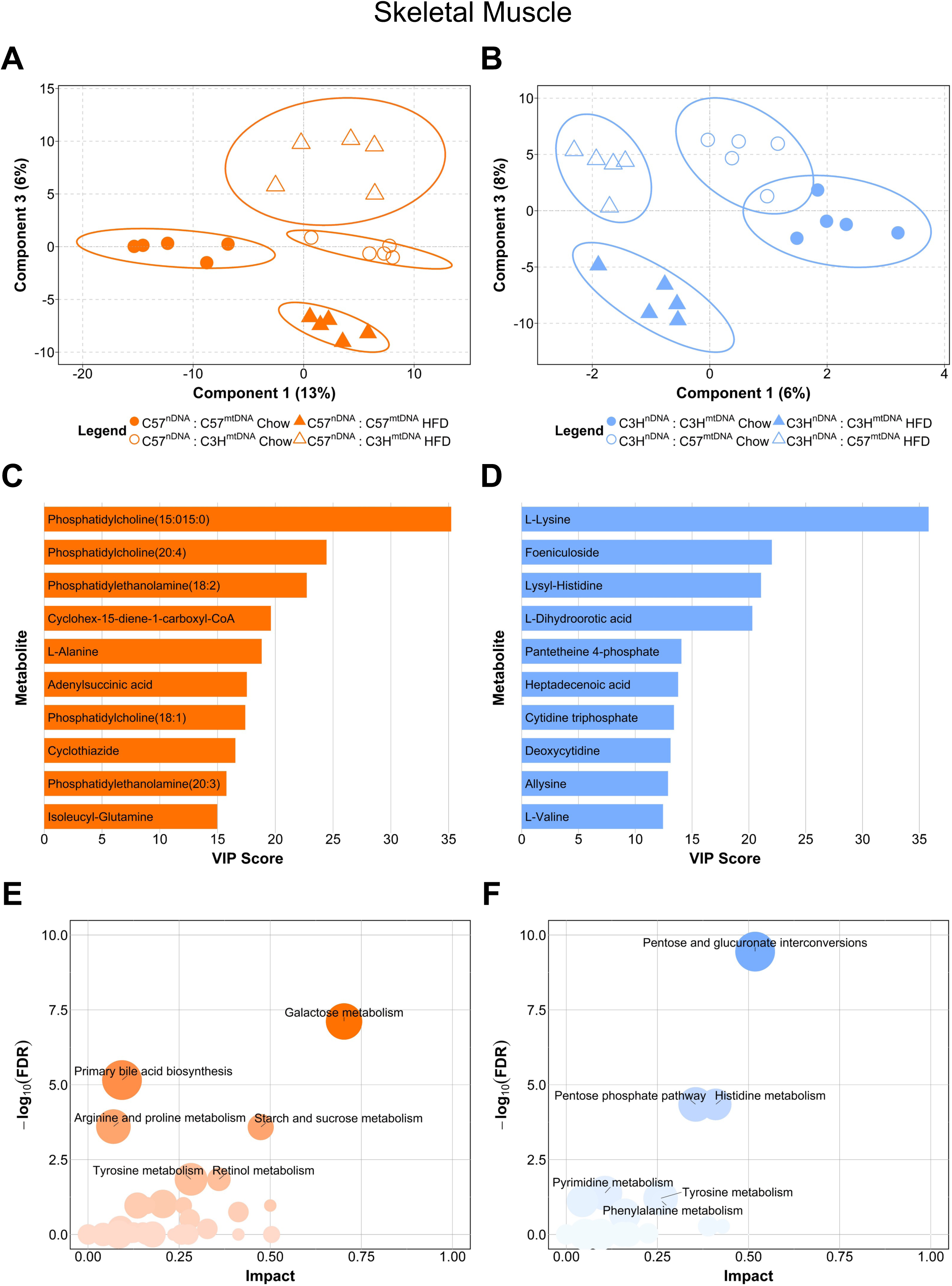
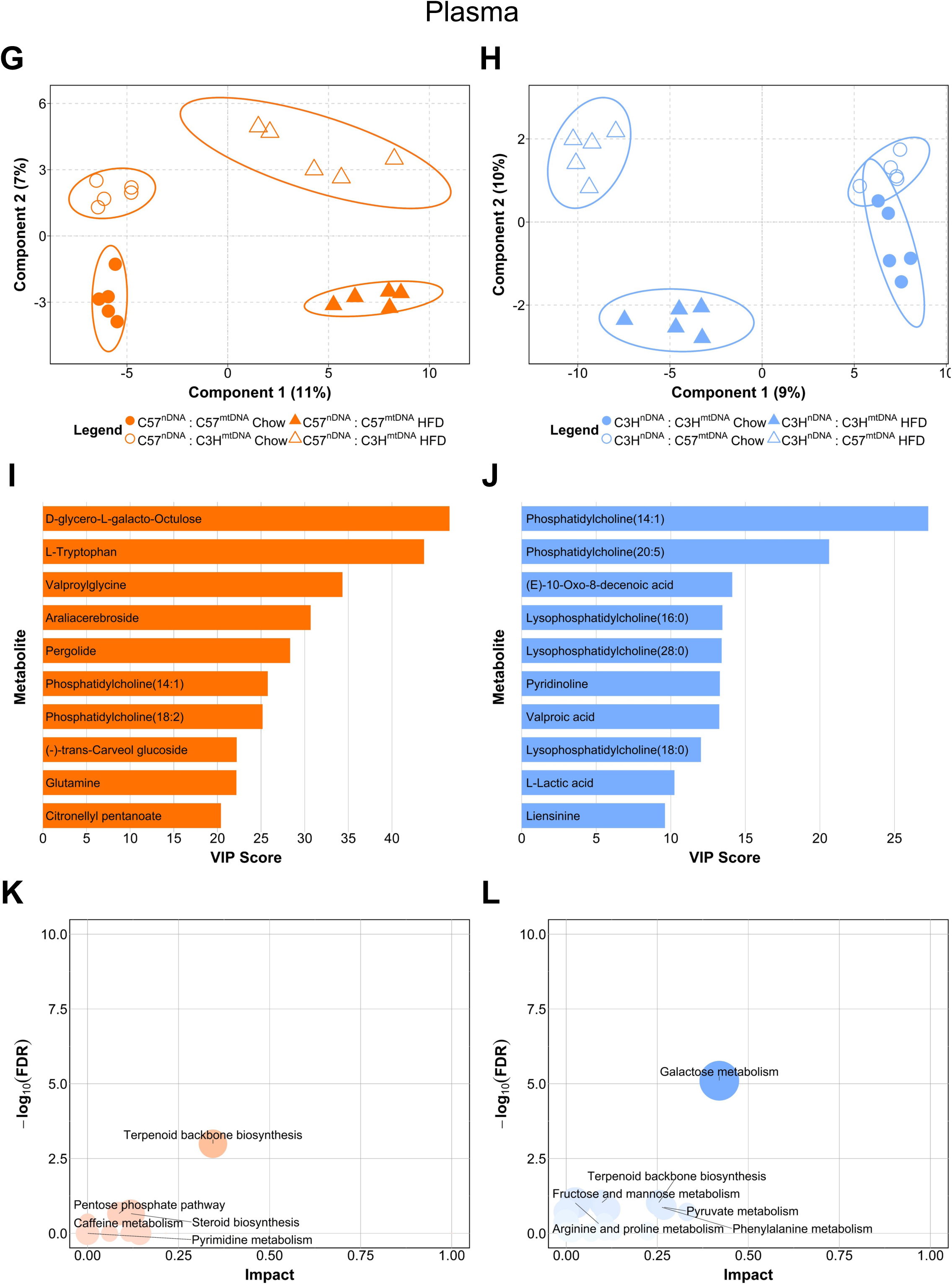
Sparse Partial Least Squares-Discriminant Analysis of Skeletal Muscle and Plasma in mice with C57^nDNA^ and C3H^nDNA^. **(A)** PLS-DA biplot of the first and third principal components of Skeletal Muscle from mice possessing C57^nDNA^ (C57^nDNA^:C57^mtDNA^ WT and C57^nDNA^:C3H^mtDNA^ MNX) on Control (‘Chow’) and High-Fat Diet (HFD). **(B)** PLS-DA biplot of the first and third principal components of Skeletal Muscle from mice possessing C3H^nDNA^ (C3H^nDNA^:C3H^mtDNA^ WT and C3H^nDNA^:C57^mtDNA^ MNX) on Control (‘Chow’) and HFD. **(C)** The top 10 skeletal muscle metabolites separating Control Diet and HFD mice possessing C57^nDNA^ (C57^nDNA^:C57^mtDNA^ WT and C57^nDNA^:C3H^mtDNA^ MNX). Metabolites are ranked according to Variable Importance in Projection (VIP) scores along the third principal component. **(D)** The top 10 skeletal muscle metabolites separating Control Diet and HFD mice possessing C3H^nDNA^ (C3H^nDNA^:C3H^mtDNA^ WT and C3H^nDNA^:C57^mtDNA^ MNX). Metabolites are ranked according to Variable Importance in Projection (VIP) Scores along the third principal component. **(E)** Bubble plot of KEGG-based pathway analysis of all Skeletal Muscle C57^nDNA^ metabolites from (A) with VIP Score > 0 along the third principal component. Pathway terms are plotted along –log_10_(FDR) and impact factor. Bubble size represents number of Hits. KEGG pathway terms with FDR < 0.05 are annotated. **(F)** Bubble plot of KEGG-based pathway analysis of all Skeletal Muscle C3H^nDNA^ metabolites from (C) with VIP Score > 0 along the third principal component. Pathway terms are plotted along –log_10_(FDR) and impact factor. Bubble size represents number of Hits. KEGG pathway terms with FDR < 0.05 are annotated. **(G)** PLS-DA biplot of the first and second principal components of Plasma from mice possessing C57^nDNA^ (C57^nDNA^:C57^mtDNA^ WT and C57^nDNA^:C3H^mtDNA^ MNX) on Control (‘Chow’) and HFD. **(H)** PLS-DA biplot of the first and second principal components of Plasma from mice possessing C3H^nDNA^ (C3H^nDNA^:C3H^mtDNA^ WT and C3H^nDNA^:C57^mtDNA^ MNX) on Control (‘Chow’) and HFD. **(I)** The top 10 Plasma metabolites separating Control Diet and HFD mice possessing C57^nDNA^ (C57^nDNA^:C57^mtDNA^ WT and C57^nDNA^:C3H^mtDNA^ MNX). Metabolites are ranked according to Variable Importance in Projection (VIP) scores along the second principal component. **(J)** The top 10 Plasma metabolites separating Control Diet and HFD mice possessing C3H^nDNA^ (C3H^nDNA^:C3H^mtDNA^ WT and C3H^nDNA^:C57^mtDNA^ MNX). Metabolites are ranked according to Variable Importance in Projection (VIP) Scores along the second principal component. **(K)** Bubble plot of KEGG-based pathway analysis of all Plasma C57^nDNA^ metabolites from (A) with VIP Score > 0 along the second principal component. Pathway terms are plotted along –log_10_(FDR) and impact factor. Bubble size represents number of Hits. KEGG pathway terms with FDR < 0.05 are annotated. **(L)** Bubble plot of KEGG-based pathway analysis of all Plasma C3H^nDNA^ metabolites from (C) with VIP Score > 0 along the second principal component. Pathway terms are plotted along – log_10_(FDR) and impact factor. Bubble size represents number of Hits. KEGG pathway terms with FDR < 0.05 are annotated.

The top 10 skeletal muscle metabolites in mice possessing C57^nDNA^ according to Variable Importance of Projection (VIP) scores in the third PComp included several species of phosphatidylethanolamines (PEs) including PE(18:2) and PE(20:3); phosphatidylcholines (PCs) including PC(15:015:0), PC(20:4), and PC(18:1); L-alanine; and adenylsuccinic acid (**Figure 2C**). In the skeletal muscle of mice possessing C3H^nDNA^, several unique amino acids including L-lysine, lysyl-histidine, allysine, and L-valine, as well as heptadecenoic acid and pantetheine 4-phosphate comprised the top 10 metabolites by VIP score, which explained variance along the third PComp (**Figure 2D**).

In mice possessing C57^nDNA^, the three most significant pathway terms identified from these metabolites included ‘Galactose metabolism’ (mmu00052; FDR = 8.14 × 10^-4^), ‘Primary bile acid biosynthesis’ (mmu00120; FDR = 5.76 × 10^-3^), and ‘Starch and sucrose metabolism’ (mmu00500; FDR = 2.77 × 10^-^2, **Figure 2E**). In mice possessing C3H^nDNA^, the three most significant pathway terms identified from these metabolites included ‘Pentose and glucuronate interconversions’ (mmu00040; FDR = 7.90 × 10^-5^), ‘Pentose phosphate pathway’ (mmu00030; FDR = 1.30 × 10^-2^), and ‘Histidine metabolism’ (mmu00340; FDR = 1.30 × 10^-2^, **Figure 2F**). Interestingly, it appeared that sugar metabolism pathways separated control and HFD groups in mice possessing C57^nDNA^, whereas pentose-associated pathways characterized the separation of diet groups in mice possessing C3H^nDNA^.

#### Plasma

In the plasma of C57^nDNA^:C57^mtDNA^ WT and C57^nDNA^:C3H^mtDNA^ MNX mice, there was clear separation of all diet and strain groups along the first and second PComps. The second PComp displayed the greatest separation between HFD and control groups (**Figure 2G**). There was no separation of control diet groups in C3H^nDNA^:C3H^mtDNA^ WT and C3H^nDNA^:C57^mtDNA^ MNX plasma. Similar to mice possessing C57^nDNA^, the second PComp displayed the greatest separation between control and HFD groups (**Figure 2H**).

In the plasma of mice possessing C57^nDNA^, several amino acids including L-tryptophan, valproylglycine, L-glutamine, which were not found in skeletal muscle, as well as PCs including PC(14:1) and PC(18:2), comprised the top 10 metabolites by VIP score, driving control and HFD group separation along the second PComp (**Figure 2I**). Interestingly, PCs with chain length 18 were found in both skeletal muscle and plasma sPLS-DA analyses of mice harboring C57^nDNA^. The top 10 plasma metabolites by VIP score in mice possessing C3H^nDNA^, representing the variance along the second PComp, includes PCs such as PC(14:1) and PC(20:5); lysophosphatidylcholines (LysoPCs) including LysoPC(16:0), LysoPC(28:0), and LysoPC(18:0); and L-lactic acid (**Figure 2J**).

In mice possessing C57^nDNA^, the three most significant pathway terms identified from these metabolites were ‘Terpenoid backbone biosynthesis’ (mmu00900; FDR = 5.0134 × 10^-2^), ‘Pentose phosphate pathway’ (mmu00030; FDR = 5.1787 × 10^-1^), and ‘Steroid biosynthesis’ (mmu00100; FDR = 5.1787 × 10^-1^, **Figure 2K**). In mice possessing C3H^nDNA^, the three most significant pathway terms identified from these metabolites included ‘Galactose metabolism’ (mmu00052; FDR = 6.0826 × 10^-3^), ‘Terpenoid backbone biosynthesis’ (mmu00900; FDR = 3.591 × 10^-1^), and ‘Fructose and mannose metabolism’ (mmu00051; FDR = 3.591 × 10^-1^, **Figure 2L**).

In plasma, the exploratory sPLS-DA analyses revealed that steroid synthesis and pentose phosphate pathways were enriched in mice harboring C57^nDNA^; in mice harboring C3H^nDNA^, galactose and fructose metabolic pathways were identified. Interestingly, these trends were reversed in skeletal muscle. To further resolve differential responses to HFD feeding in WT and MNX mice, differential abundance analysis was performed in nDNA-matched mice fed HFD against controls.

### Differential Abundance Analysis Reveals Strain-Specific Response to HFD

#### Effect of High-Fat Diet on Skeletal Muscle of C57^nDNA^:C57^mtDNA^ WT Mice

In C57^nDNA^:C57^mtDNA^ WT mice, HFD significantly altered skeletal muscle glycerophospholipid and glucose profiles compared to controls. Downregulation of LysoPC(16:0) and dimethylphosphatidylethanolamine (PE-NME2) PE-NME2(22:6) (*P*-adj < 0.05), may reflect greater dysregulation of lipid beta-oxidation ^100^. Additionally, D-Glucose was downregulated (*P*-adj < 0.05), which may be indicative of lower glucose import into skeletal muscle (**Figure 3A**). To contextualize these metabolites, KEGG-based pathway enrichment analysis revealed ‘Galactose metabolism’ (mmu00052; *P*-value = 1.38 × 10^-9^), ‘Fructose and mannose metabolism’ (mmu00051; *P*-value = 1.10 × 10^-4^), and ‘Starch and sucrose metabolism’ (mmu00500; *P*-value = 3.1176 × 10^-3^) as notable pathways due to the differential regulation of keto-b-D-galactose and D-Glucose (**Figure 3B**).

**Figure 3:**
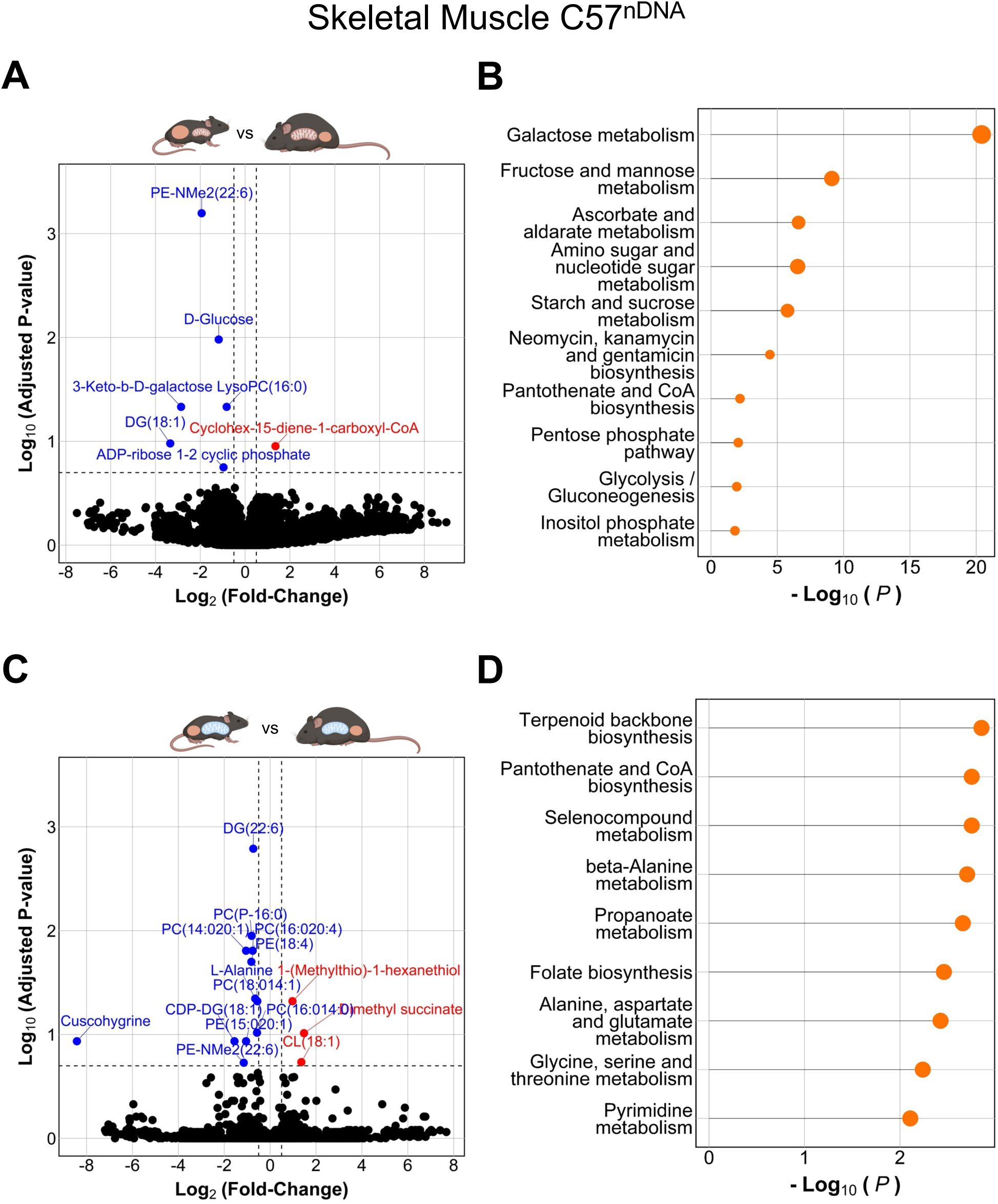
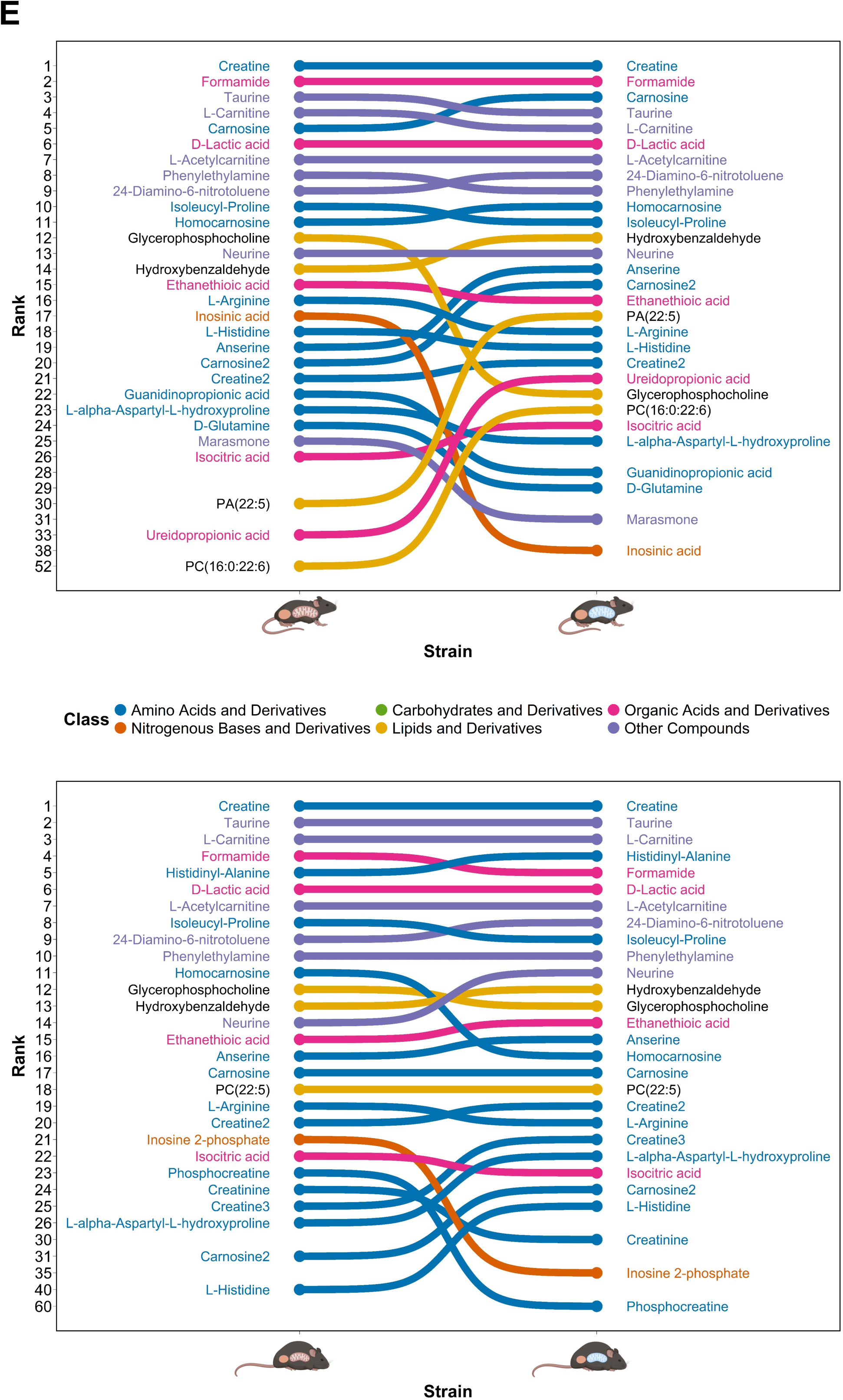
Differential Abundance Analysis of the effect of HFD vs Control in Skeletal Muscle and ranking of the top 25 Skeletal Muscle metabolites within Diet-matched mice, harboring C57^nDNA^. **(A)** Volcano plot of differentially abundant metabolites in C57^nDNA^:C57^mtDNA^ WT mice fed High-Fat Diet (HFD) relative to Control Diet. The horizontal red line represents an adjusted *P*-value of 0.2 and the vertical red line represents 1-fold change relative to Control mice. Differentially abundant metabolites meeting the adjusted *P*-value of 0.2 threshold are coloured blue if downregulated and red if upregulated. The top 10 metabolites by adjusted *P*-value are annotated. **(B)** KEGG-based pathway analysis of differentially abundant Skeletal Muscle metabolites from (A) in C57^nDNA^:C57^mtDNA^ WT mice. Pathway terms are plotted along –log_10_(*P*-value). Bubble size represents number of Hits. The top 10 pathway terms by *P*-value are displayed. **(C)** Volcano plot of differentially abundant metabolites in C57^nDNA^:C3H^mtDNA^ MNX mice fed HFD relative to Control Diet. The horizontal red line represents an adjusted *P*-value of 0.2 and the vertical red line represents 1-fold change relative to Control mice. Differentially abundant metabolites meeting the adjusted *P*-value of 0.2 threshold are coloured blue if downregulated and red if upregulated. The top 10 metabolites by adjusted *P*-value are annotated. **(D)** KEGG-based pathway analysis of differentially abundant Skeletal Muscle metabolites from (C) in C57^nDNA^:C3H^mtDNA^ MNX mice. Pathway terms are plotted along –log_10_(*P*-value). Bubble size represents number of Hits. The top 10 pathway terms by *P*-value are displayed. **(E)** Bump chart of the top 25 metabolites by mean peak intensity in **(top)** Control Diet-fed and **(bottom)** HFD-fed C57^nDNA^:C57^mtDNA^ WT (orange nucleus, orange mitochondria) and C57^nDNA^:C3H^mtDNA^ MNX (orange nucleus, blue mitochondria). The respective ranks of metabolites in the opposing nDNA-matched strain are also listed to display strain-specific changes in the most abundant metabolites within Skeletal Muscle. Metabolites are coloured by general biochemical class: Amino Acids and Derivatives (Blue), Nitrogenous Bases and Derivatives (Orange), Carbohydrates and Derivatives (Green), Lipids and Derivatives (Yellow), Organic Acids and Derivatives (Pink), and Other Compounds (Purple).

#### Effect of High-Fat Diet on Skeletal Muscle of C57^nDNA^:C3H^mtDNA^ MNX Mice

In the skeletal muscle of C57^nDNA^:C3H^mtDNA^ MNX mice, HFD significantly downregulated several species of long-chain PCs including PC(P-16:0), PC(14:020:1), PC(18:014:1), and PC(16:014:0), and long-chain PEs including PE(18:4), PE(15:020:1), and PE-NMe2(22:6) (P-adj < 0.2). PE-NME2(22:6) was downregulated in both C57^nDNA^:C57^mtDNA^ WT and C57^nDNA^:C3H^mtDNA^ MNX mice fed HFD, however, this change was larger (log_2_FC = -1.94 vs. -1.15) and more significant in the WT strain compared to C57^nDNA^:C3H^mtDNA^ MNX (*P*-adj = 6.36 × 10^-4^ vs. 1.87 × 10^-1^). This may indicate a greater lipid beta-oxidation dysregulation in mice with C57^mtDNA^ background relative to C3H^mtDNA^ ^100^. There were no significant changes to short- or medium-chain glycerophospholipids, although this may be expected as long-chain glycerophospholipids are the most abundant within the glycerophospholipid class ^101^. L-alanine and diacylglyercols (DGs) such as DG(22:6) and CDP-DG(18:1) were also downregulated (*P*-adj < 0.2). Interestingly, cardiolipin (CL) CL(18:1) was upregulated (*P*-adj < 0.2) in C57nDNA:C3HmtDNA MNX mice fed HFD but not C57^nDNA^:C57^mtDNA^ WT mice suggesting that mtDNA genotype may have influenced the abundance of CLs with chain length 18, which are mitochondrial membrane lipids ^102^ (**Figure 3C**). Notable pathways enriched in C57^nDNA^:C3H^mtDNA^ MNX mice included ‘Terpenoid backbone biosynthesis’ (mmu00900; *P*-value = 5.76 × 10^-2^), ‘pantothenate and coenzyme A (CoA) biosynthesis’ (mmu00770; *P*-value = 6.39 × 10^-2^), and ‘beta-Alanine metabolism’ (mmu00410; 6.70 × 10^-2^, **Figure 3D**). Altered regulation of alanine metabolism was also reflected in the enrichment of the ‘Alanine, aspartate and glutamate metabolism’ KEGG term (mmu00250; *P*-value = 8.8494 × 10^-2^).

#### Changes in the Most Abundant Skeletal Muscle Metabolites in C57 Nuclear Backgrounds

The top 25 metabolites from diet-matched C57^nDNA^:C57^mtDNA^ WT and C57^nDNA^:C3H^mtDNA^ MNX mice were sorted and ranked by descending mean intensity (**Figure 3E**). Notable rank changes were defined as those > 2 levels in the strain with opposing mtDNA. On control diet, glycerophosphocholine, isosinic acid, guanidopropioninc acid, D-glutamine, and marasmone were ranked lower, and anserine, carnosine, phosphatidic acid (PA) PA(22:5), ureidopropionic acid, and PC(16:0:22:6) were ranked higher in C57^nDNA^:C3H^mtDNA^ MNX compared to C57^nDNA^:C57^mtDNA^ WT mice (**Figure 3E top**). On HFD, homocarnosine, inosine 2-phosphate, phosphocreatine, and creatinine were ranked lower, and neurine, creatine, L-alpha-aspartyl-L-hydroxyproline, carnosine, and L-histidine were ranked higher in C57^nDNA^:C3H^mtDNA^ MNX compared to C57^nDNA^:C57^mtDNA^ WT mice (**Figure 3E, bottom**). Interestingly, glycerophosphocholine is a metabolic cursor to various glycerophospholipids including PC, PE, and LysoPC ^103,104^, which were differentially abundant in the diet comparisons of mice harboring C57^nDNA^. Further, PCs with chain length 16 were also found to be downregulated in C57^nDNA^:C3H^mtDNA^ MNX mice fed HFD relative to controls.

#### Effect of High-Fat Diet on Skeletal Muscle of C3H^nDNA^:C3H^mtDNA^ WT Mice

In C3H^nDNA^:C3H^mtDNA^ WT mice, HFD significantly altered the skeletal muscle glycerophospholipid profiles compared to controls. In particular, long-chain PCs including PC(P-16:0) and PC(18:1) were significantly downregulated (*P*-adj < 0.2) (**Figure 4A**). Downregulation of 16 and 18 chain length PCs was also found in HFD-fed C57^nDNA^:C3H^mtDNA^ MNX mice compared to controls, indicating mtDNA background-based modulation of these metabolites. Additionally, L-alanine was significantly downregulated in HFD-fed C3H^nDNA^:C3H^mtDNA^ WT mice (*P*-adj < 0.2) and was also downregulated in C57^nDNA^:C3H^mtDNA^ MNX skeletal muscle. Pathway enrichment analysis of these metabolites noted significant enrichment of ‘Pantothenate and CoA biosynthesis’ (mmu00770; *P*-value = 3.8805 × 10^-2^) and ‘beta-Alanine metabolism’ (mmu00410; *P*-value = 4.07 × 10^-2^, **Figure 4B**).

**Figure 4:**
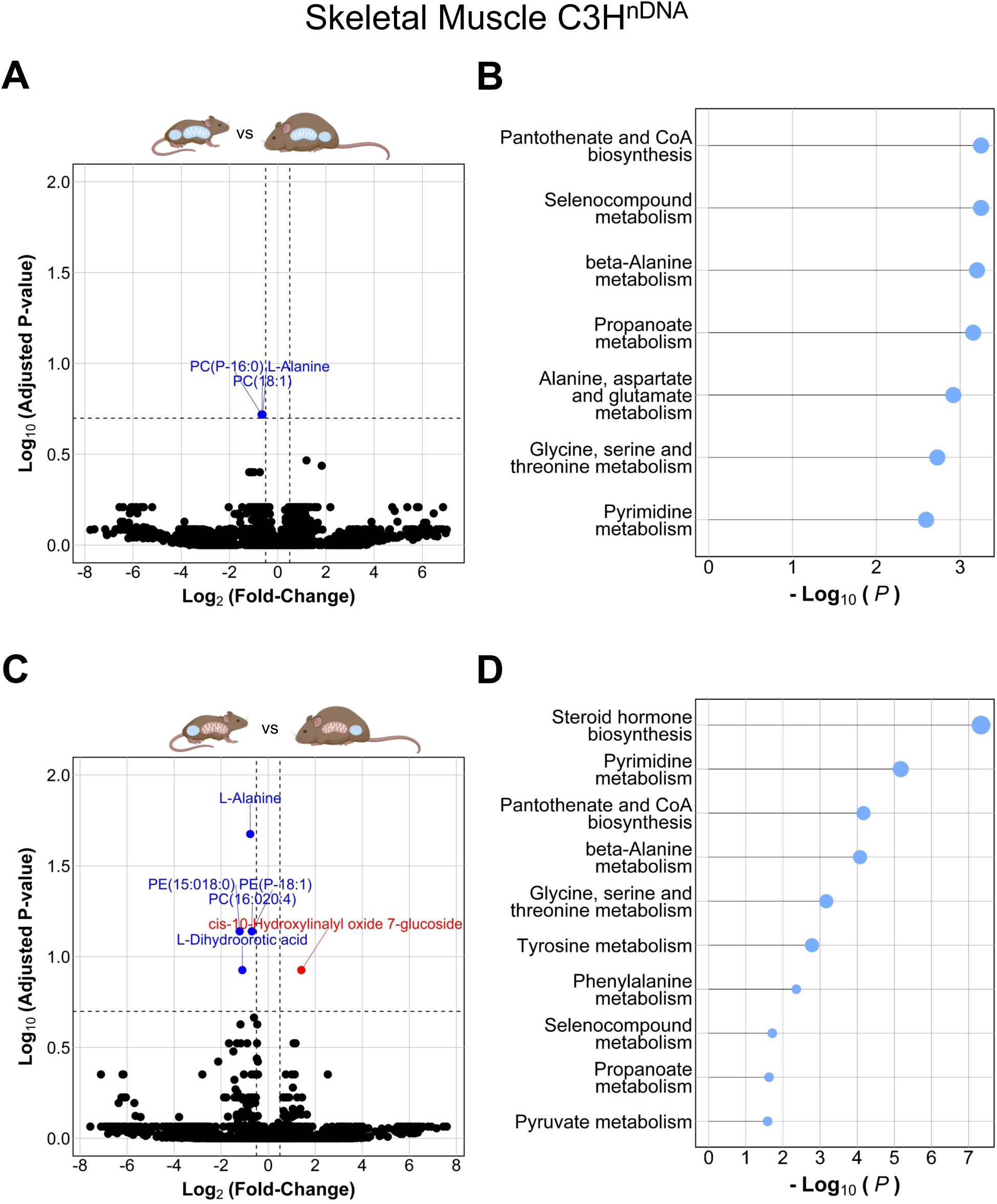
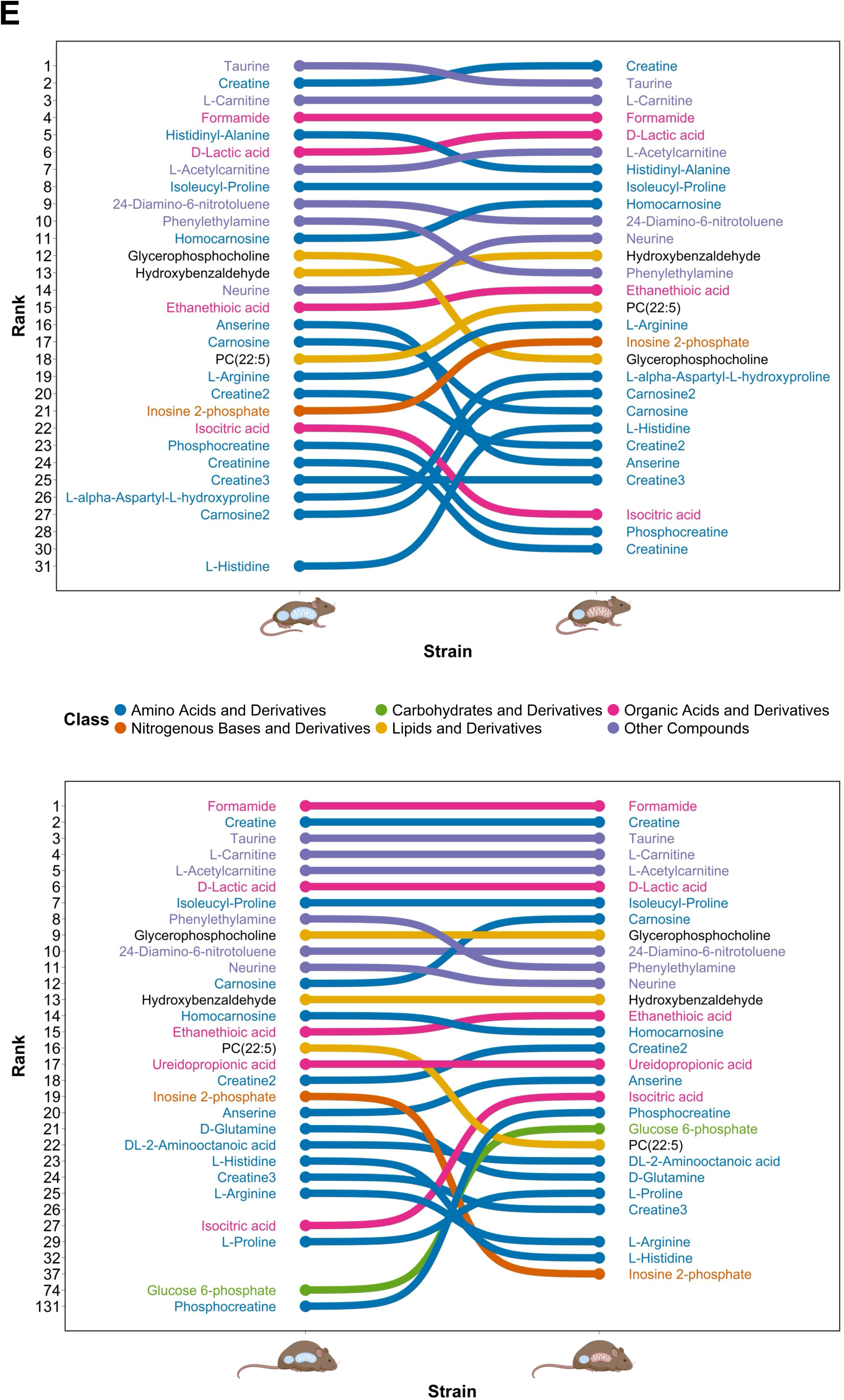
Differential Abundance Analysis of the effect of HFD vs Control in Skeletal Muscle and ranking of the top 25 Skeletal Muscle metabolites within Diet-matched mice, harboring C3H^nDNA^. **(A)** Volcano plot of differentially abundant metabolites in C3H^nDNA^:C3H^mtDNA^ WT mice fed High-Fat Diet (HFD) relative to Control Diet. The horizontal red line represents an adjusted *P*-value of 0.2 and the vertical red line represents 1-fold change relative to Control mice. Differentially abundant metabolites meeting the adjusted *P*-value of 0.2 threshold are coloured blue if downregulated and red if upregulated. The top 10 metabolites by adjusted *P*-value are annotated. **(B)** KEGG-based pathway analysis of differentially abundant Skeletal Muscle metabolites from (A) in C3H^nDNA^:C3H^mtDNA^ WT mice. Pathway terms are plotted along –log_10_(*P*-value). Bubble size represents number of Hits. The top 10 pathway terms by *P*-value are displayed. **(C)** Volcano plot of differentially abundant metabolites in C3H^nDNA^:C57^mtDNA^ MNX mice fed HFD relative to Control Diet. The horizontal red line represents an adjusted *P*-value of 0.2 and the vertical red line represents 1-fold change relative to Control mice. Differentially abundant metabolites meeting the adjusted *P*-value of 0.2 threshold are coloured blue if downregulated and red if upregulated. The top 10 metabolites by adjusted *P*-value are annotated. **(D)** KEGG-based pathway analysis of differentially abundant Skeletal Muscle metabolites from (C) in C3H^nDNA^:C57^mtDNA^ MNX mice. Pathway terms are plotted along –log_10_(*P*-value). Bubble size represents number of Hits. The top 10 pathway terms by *P*-value are displayed. **(E)** Bump chart of the top 25 metabolites by mean peak intensity in **(top)** Control Diet-fed and **(bottom)** HFD-fed C3H^nDNA^:C3H^mtDNA^ WT (blue nucleus, blue mitochondria) and C3H^nDNA^:C57^mtDNA^ MNX (blue nucleus, orange mitochondria). The respective ranks of metabolites in the opposing nDNA-matched strain are also listed to display strain-specific changes in the most abundant metabolites within Skeletal Muscle. Metabolites are coloured by general biochemical class: Amino Acids and Derivatives (Blue), Nitrogenous Bases and Derivatives (Orange), Carbohydrates and Derivatives (Green), Lipids and Derivatives (Yellow), Organic Acids and Derivatives (Pink), and Other Compounds (Purple).

#### Effect of High-Fat Diet on Skeletal Muscle of C3H^nDNA^:C57^mtDNA^ MNX Mice

In C3H^nDNA^:C57^mtDNA^ MNX mice, HFD significantly altered long-chain glycerophospholipids compared to controls. In particular, PC(16:020:4), PE(15:018:0), and PE(P-18:1) were significantly downregulated (*P*-adj < 0.1). As seen in mice possessing C57^nDNA^, no short- or medium-chain glycerophospholipids were altered. Interestingly, downregulated glycerophospholipids in HFD-fed C3H^nDNA^:C57^mtDNA^ MNX mice largely comprised of PEs, whereas downregulated glycerophospholipids in HFD-fed C3H^nDNA^:C3H^mtDNA^ WT mice were entirely PCs; this may indicate differential metabolism of dietary fat in a genotype-specific manner. L-alanine was downregulated in HFD-fed C57^nDNA^:C3H^mtDNA^ MNX, C3H^nDNA^:C3H^mtDNA^ WT, and C3H^nDNA^:C57^mtDNA^ MNX mice compared to their respective controls (*P*-adj < 0.2); it is unclear if L-alanine is influenced by a particular genetic background, but we note consistent downregulation in mice possessing the C3H^mtDNA^ background, whereas alanine is not differentially abundant in C57^nDNA^:C57^mtDNA^ WT skeletal muscle (**Figure 4C**). Notable pathways enriched in C3H^nDNA^:C57^mtDNA^ MNX skeletal muscle include ‘Steroid hormone biosynthesis’ (mmu00140; *P*-value = 6.52 × 10^-4^), ‘Pantothenate and CoA biosynthesis’ (mmu00770; *P*-value = 1.55 × 10^-2^), and ‘beta-Alanine metabolism’ (mmu00410; *P*-value = 1.70 × 10^-2^, **Figure 4D**).

#### Changes in the Most Abundant Skeletal Muscle Metabolites in C3H Nuclear Backgrounds

On control diet, glycerophosphocholine, anserine, carnosine, creatine, and isocitric acid were ranked lower, and PC(22:5), L-arginine, inosine 2-phosphate, L-alpha-aspartyl-L-hydroxyproline, carnosine, and L-histidine were ranked higher in C3H^nDNA^:C57^mtDNA^ MNX mice compared to C3H^nDNA^:C3H^mtDNA^ WT (**Figure 4E, top**). On HFD, phenylethylamine, PC(22:5), inosine 2-phosphate, D-glutamine, L-histidine, and L-arginine were ranked lower, and carnosine, isocitric acid, L-proline, glucose 6-phosphate, and phosphocreatine were ranked higher in C3H^nDNA^:C57^mtDNA^ MNX mice compared to C3H^nDNA^:C3H^mtDNA^ WT (**Figure 4E, bottom**).

#### Effect of High-Fat Diet on Plasma of C57^nDNA^:C57^mtDNA^ WT Mice

In the plasma of C57^nDNA^:C57^mtDNA^ WT mice, HFD-feeding significantly altered 275 metabolites in HFD-fed mice relative to controls. The most significantly regulated metabolites included various glycerophospholipids and storage lipids, including phosphatidylglycerolphosphate (PGP) PGP(18:3), PE(22:4), PA(18:1), DG(18:3), DG(20:4), and cytidine diphosphate (CDP)-DG(22:6), which were significantly downregulated, and PC(15:018:3) and CDP-DG(20:4) which were significantly upregulated (*P*-adj < 0.2, **Figure 5A**). Interestingly, the diversity of plasma glycerophospholipids was greater than that found in skeletal muscle, yet the most differentially abundant glycerophospholipids were long-chain fatty acids, similar to those found in skeletal muscle. ‘Steroid biosynthesis’ (mmu00100; *P*-value = 2.17 × 10^-4^), ‘Galactose metabolism’ (mmu00052; *P*-value = 8.04 × 10^-4^), and ‘Tyrosine metabolism’ (mmu00350; *P*-value = 1.19 × 10^-3^) pathways were notably significantly enriched (**Figure 5B**).

**Figure 5:**
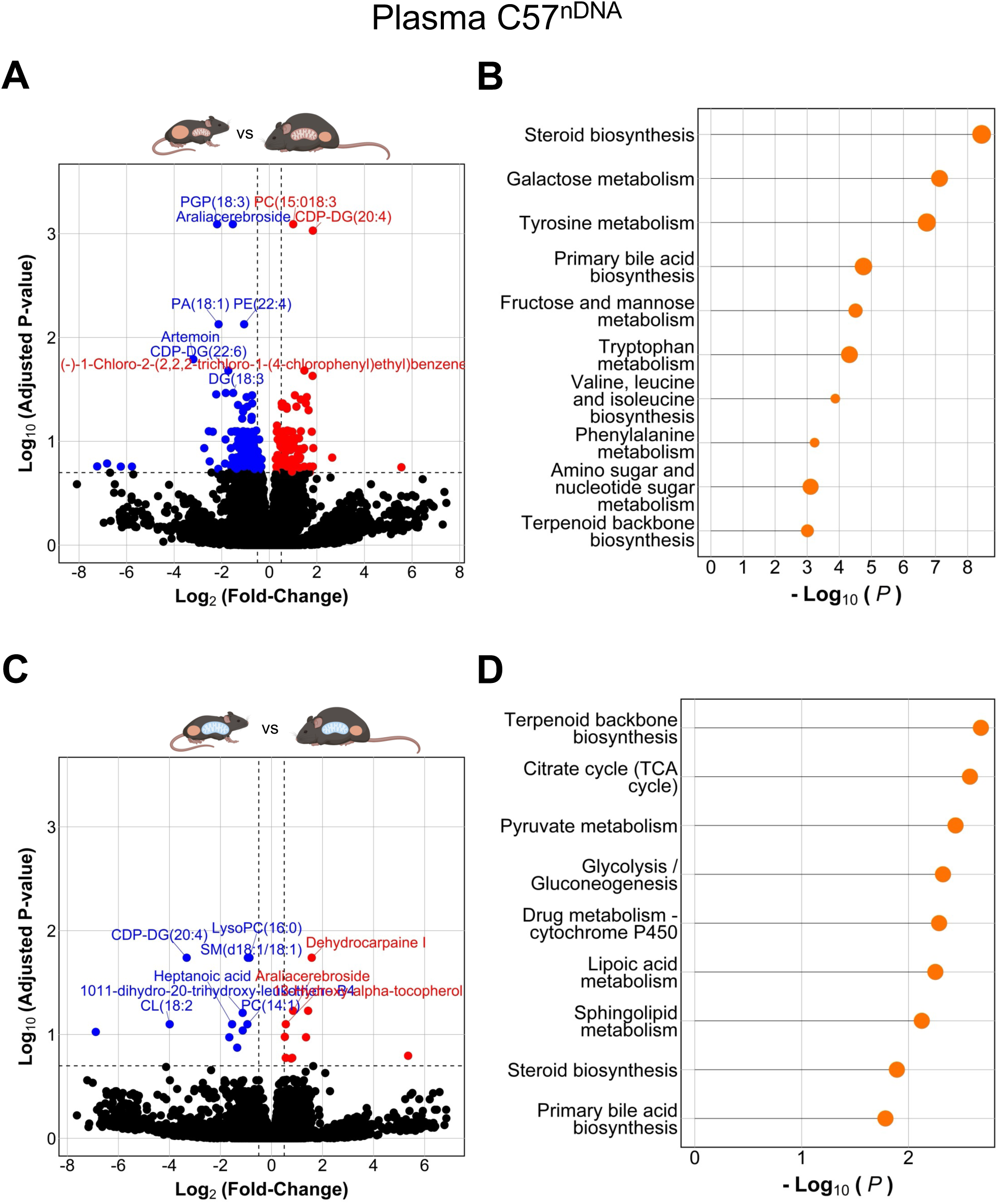
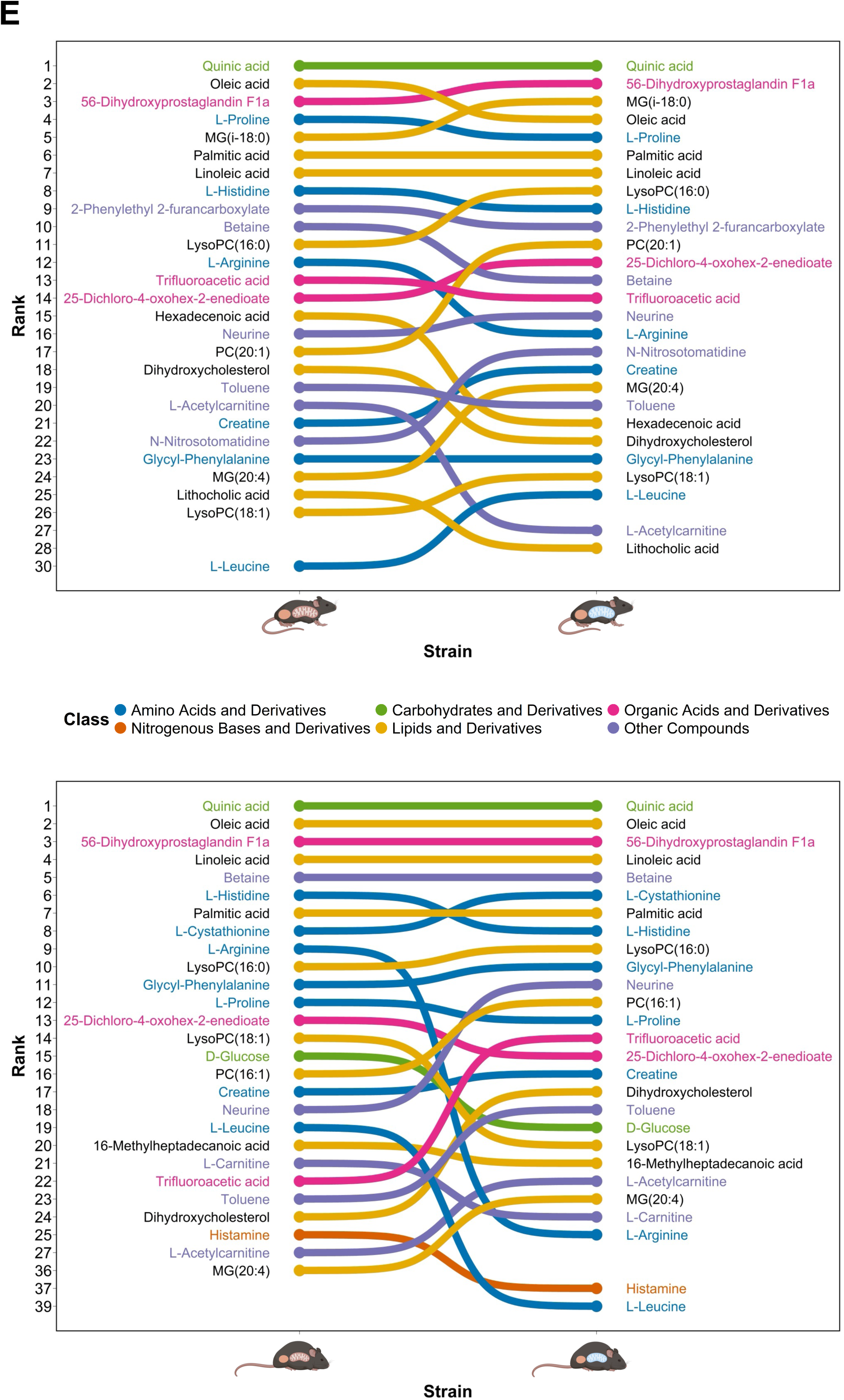
Differential Abundance Analysis of the effect of HFD vs Control in Plasma and ranking of the top 25 Plasma metabolites within Diet-matched mice, harboring C57^nDNA^. **(A)** Volcano plot of differentially abundant metabolites in C57^nDNA^:C57^mtDNA^ WT mice fed High-Fat Diet (HFD) relative to Control Diet. The horizontal red line represents an adjusted *P*-value of 0.2 and the vertical red line represents 1-fold change relative to Control mice. Differentially abundant metabolites meeting the adjusted *P*-value of 0.2 threshold are coloured blue if downregulated and red if upregulated. The top 10 metabolites by adjusted *P*-value are annotated. **(B)** KEGG-based pathway analysis of differentially abundant Plasma metabolites from (A) in C57^nDNA^:C57^mtDNA^ WT mice. Pathway terms are plotted along –log_10_(*P*-value). Bubble size represents number of Hits. The top 10 pathway terms by *P*-value are displayed. **(C)** Volcano plot of differentially abundant metabolites in C57^nDNA^:C3H^mtDNA^ MNX mice fed HFD relative to Control Diet. The horizontal red line represents an adjusted *P*-value of 0.2 and the vertical red line represents 1-fold change relative to Control mice. Differentially abundant metabolites meeting the adjusted *P*-value of 0.2 threshold are coloured blue if downregulated and red if upregulated. The top 10 metabolites by adjusted *P*-value are annotated. **(D)** KEGG-based pathway analysis of differentially abundant Plasma metabolites from (C) in C57^nDNA^:C3H^mtDNA^ MNX mice. Pathway terms are plotted along –log_10_(*P*-value). Bubble size represents number of Hits. The top 10 pathway terms by *P*-value are displayed. **(E)** Bump chart of the top 25 metabolites by mean peak intensity in **(top)** Control Diet-fed and **(bottom)** HFD-fed C57^nDNA^:C57^mtDNA^ WT (orange nucleus, orange mitochondria) and C57^nDNA^:C3H^mtDNA^ MNX (orange nucleus, blue mitochondria). The respective ranks of metabolites in the opposing nDNA-matched strain are also listed to display strain-specific changes in the most abundant metabolites within Plasma. Metabolites are coloured by general biochemical class: Amino Acids and Derivatives (Blue), Nitrogenous Bases and Derivatives (Orange), Carbohydrates and Derivatives (Green), Lipids and Derivatives (Yellow), Organic Acids and Derivatives (Pink), and Other Compounds (Purple).

#### Effect of High-Fat Diet on Plasma of C57^nDNA^:C3H^mtDNA^ MNX Mice

In C57^nDNA^:C3H^mtDNA^ MNX mice, HFD-feeding significantly altered 21 metabolites compared to controls. Notably, sphingolipids such as sphingomyelin (SM) SM(d18:1/18:1), glycerophospholipids such as CDP-DG(20:4), LysoPC(16:0), CL(18:2), and PC(14:1), and short-chain fatty acids such as heptanoic acid were the most significantly downregulated metabolites (*P*-adj < 0.2, **Figure 5C**). The chain length of circulating glycerophospholipids remains large relative to skeletal muscle; interestingly, LysoPC(16:0), PC(14:1), and CLs with chain length 18 were found in both *C57^nDNA^:C3H^mtDNA^* skeletal muscle and plasma, which may indicate similar regulation across tissues in mice possessing C3H^mtDNA^. Significantly enriched pathways in C57^nDNA^:C3H^mtDNA^ MNX plasma included ‘Terpenoid backbone biosynthesis’ (mmu00900; *P*-value = 6.88 × 10^-2^), ‘Citrate cycle (TCA cycle)’ (mmu00020; *P*-value = 7.62 × 10^-2^), and ‘Pyruvate metabolism’ (mmu00620; *P*-value = 8.72 × 10^-2^, **Figure 5D**).

#### Changes in the Most Abundant Plasma Metabolites in C57 Nuclear Backgrounds

On control diet, betaine, L-arginine, hexadecenoic acid, dihydroxycholesterol, L-acetylcarnitine, and lithocholic acid were ranked lower, and LysoPC(16:0), PC(20:1), creatine, N-nitrosotomatidine, monoglyceride (MG) MG(20:4), and L-leucine were ranked higher, in C57^nDNA^:C3H^mtDNA^ MNX mice compared to C57^nDNA^:C57^mtDNA^ WT (**Figure 5E, top**). On HFD, L-arginine, LysoPC(18:1), D-glucose, L-leucine, L-carnitine, and histamine were ranked lower, and PC(16:1), neurine, trifluoroacetic acid, toluene, dihydroxycholesterol, L-acetylcarnitine, and MG(20:4) were ranked higher in C57^nDNA^:C3H^mtDNA^ MNX mice compared to C57^nDNA^:C57^mtDNA^ WT (**Figure 5E, bottom**). LysoPCs with chain lengths of 16 and 18, and various long-chain PCs were highly abundant in both control diet- and HFD-fed mice and were differentially regulated, which may be indicative of changes to circulating glycerophospholipid precursor abundance (as LysoPC is a precursor to PC); circulating long-chain LysoPCs and PCs are known to be associated with altered insulin sensitivity ^105,106^, and this relationship may be modulated by distinct mtDNA backgrounds.

#### Effect of High-Fat Diet on Plasma of C3H^nDNA^:C3H^mtDNA^ WT Mice

In C3H^nDNA^:C3H^mtDNA^ WT plasma, HFD significantly altered the abundance of 131 metabolites in HFD-fed mice relative to control. Notably, CL(i-16:0/a-17:0/18:2), CL(18:2), PC(18:2), and 2-hydroxybenzaldehyde were significantly downregulated, and phosphatidylglycerol (PG) PG(20:5) was significantly upregulated (*P*-adj < 0.2, **Figure 6A**). Highly significant differential abundance of chain length 18 CLs was found in C3H^nDNA^:C3H^mtDNA^ WT plasma and C57^nDNA^:C3H^mtDNA^ MNX skeletal muscle and plasma, which may indicate C3H^mtDNA^-specific regulation of these mitochondrial glycerophospholipids. ‘Fructose and mannose metabolism’ (mmu00051; *P*-value = 4.63 × 10^-3^), ‘Steroid biosynthesis’ (mmu00100; *P*-value = 4.77 × 10^-3^), and ‘Arginine biosynthesis’ (mmu00220; *P*-value = 1.61 × 10^-2^) were notably significantly enriched (**Figure 6B**).

**Figure 6:**
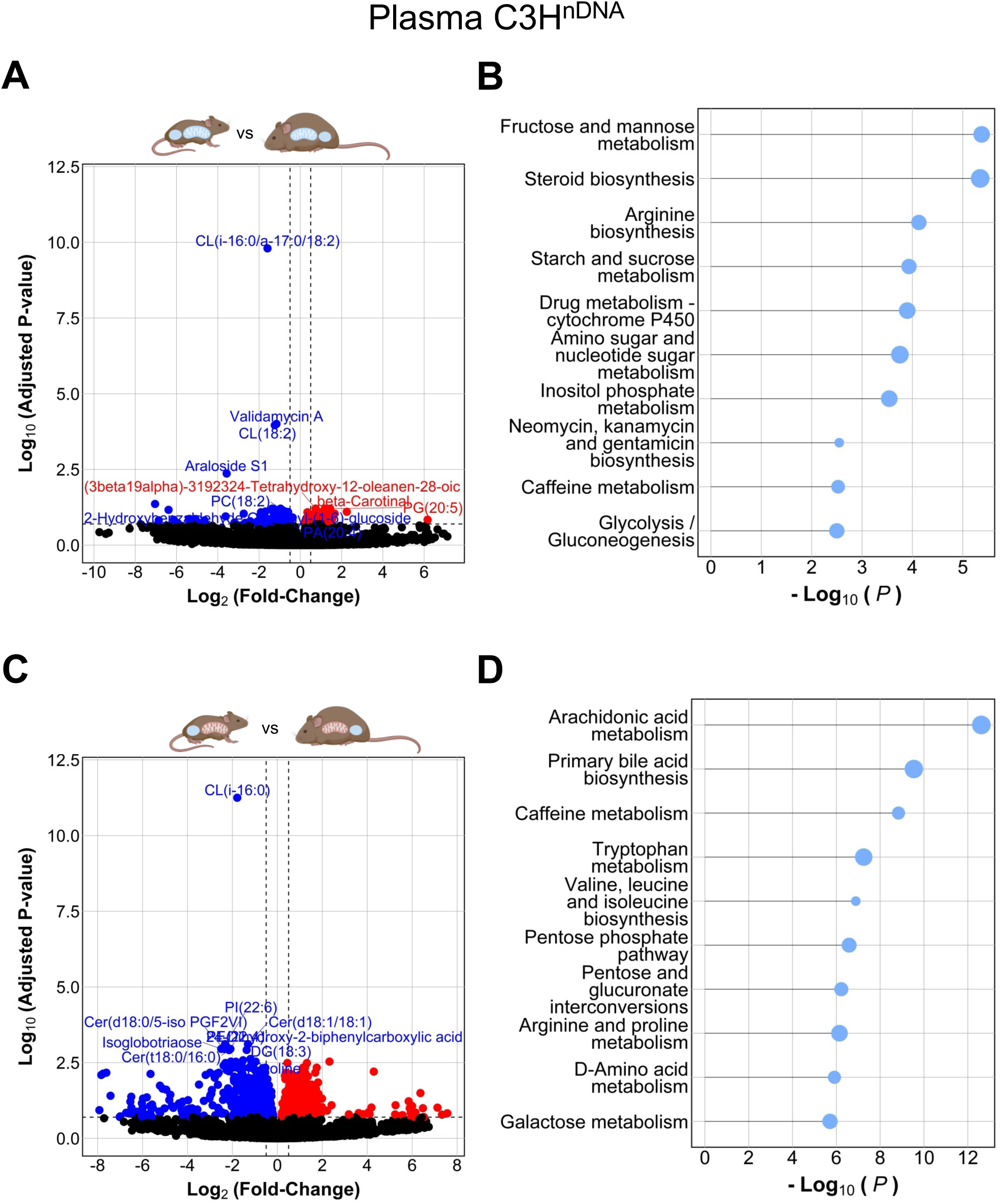
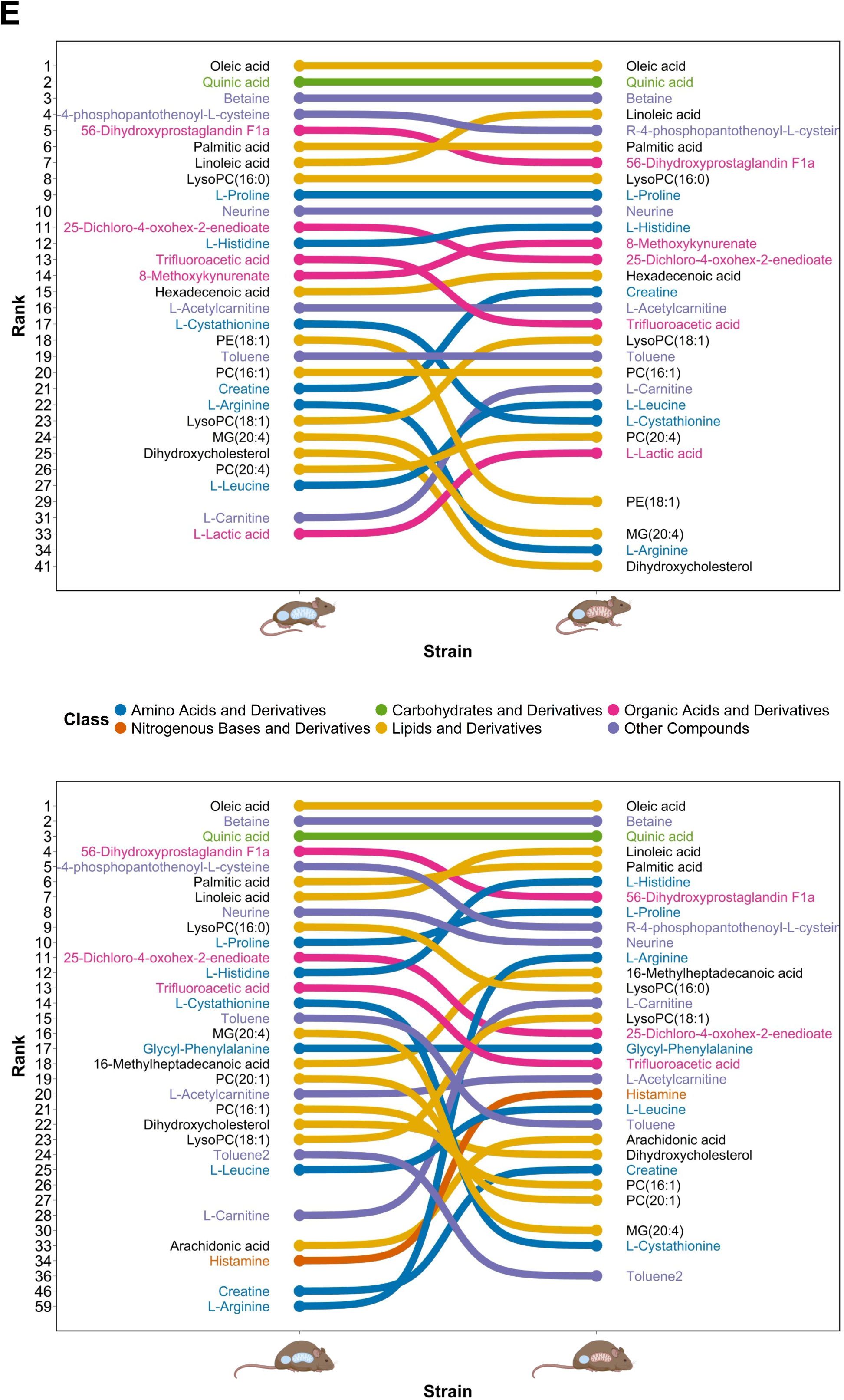
Differential Abundance Analysis of the effect of HFD vs Control in Plasma and ranking of the top 25 Plasma metabolites within Diet-matched mice, harboring C3H^nDNA^. **(A)** Volcano plot of differentially abundant metabolites in C3H^nDNA^:C3H^mtDNA^ WT mice fed High-Fat Diet (HFD) relative to Control Diet. The horizontal red line represents an adjusted *P*-value of 0.2 and the vertical red line represents 1-fold change relative to Control mice. Differentially abundant metabolites meeting the adjusted *P*-value of 0.2 threshold are coloured blue if downregulated and red if upregulated. The top 10 metabolites by adjusted *P*-value are annotated. **(B)** KEGG-based pathway analysis of differentially abundant Plasma metabolites from (A) in C3H^nDNA^:C3H^mtDNA^ WT mice. Pathway terms are plotted along –log_10_(*P*-value). Bubble size represents number of Hits. The top 10 pathway terms by *P*-value are displayed. **(C)** Volcano plot of differentially abundant metabolites in C3H^nDNA^:C57^mtDNA^ MNX mice fed HFD relative to Control Diet. The horizontal red line represents an adjusted *P*-value of 0.2 and the vertical red line represents 1-fold change relative to Control mice. Differentially abundant metabolites meeting the adjusted *P*-value of 0.2 threshold are coloured blue if downregulated and red if upregulated. The top 10 metabolites by adjusted *P*-value are annotated. **(D)** KEGG-based pathway analysis of differentially abundant Plasma metabolites from (C) in C3H^nDNA^:C57^mtDNA^ MNX mice. Pathway terms are plotted along –log_10_(*P*-value). Bubble size represents number of Hits. The top 10 pathway terms by *P*-value are displayed. **(E)** Bump chart of the top 25 metabolites by mean peak intensity in **(top)** Control Diet-fed and **(bottom)** HFD-fed C3H^nDNA^:C3H^mtDNA^ WT (blue nucleus, blue mitochondria) and C3H^nDNA^:C57^mtDNA^ MNX (blue nucleus, orange mitochondria). The respective ranks of metabolites in the opposing nDNA-matched strain are also listed to display strain-specific changes in the most abundant metabolites within Plasma. Metabolites are coloured by general biochemical class: Amino Acids and Derivatives (Blue), Nitrogenous Bases and Derivatives (Orange), Carbohydrates and Derivatives (Green), Lipids and Derivatives (Yellow), Organic Acids and Derivatives (Pink), and Other Compounds (Purple).

#### Effect of High-Fat Diet on Plasma of C3H^nDNA^:C57^mtDNA^ MNX Mice

In C3H^nDNA^:C57^mtDNA^ MNX mice, HFD significantly altered the abundance of 1,627 metabolites. Notably, CL(i-16:0), ceramides (Cer) Cer(d18:1/18:1), Cer(d18:0/5-isoPGF2VI), Cer(t18:0/16:0), phosphatidylinositol (PI) PI(22:6), PE(22:4), 24-dihydroxy-2-biphenylcarboxylic acid, isoglobotriaose, and DG(18:3) were significantly downregulated (*P*-adj < 0.2) (**Figure 6C**). Interestingly, CL(i-16:0) remained downregulated across C3H^nDNA^:C3H^mtDNA^ WT *and* C3H^nDNA^:C57^mtDNA^ MNX. Although unclear, the chain length of circulating cardiolipins may be modulated by nDNA or mtDNA background. Significantly enriched pathways in C3H^nDNA^:C57^mtDNA^ MNX plasma included ‘Arachidonic acid metabolism’ (mmu00590; *P*-value = 3.30 × 10^-6^), ‘Primary bile acid biosynthesis’ (mmu00120; *P*-value = 7.19 × 10^-5^), and ‘Tryptophan metabolism’ (mmu00380; *P*-value = 7.11 × 10^-4^, **Figure 6D**).

#### Changes in the Most Abundant Plasma Metabolites in C3H Nuclear Backgrounds

On control diet, trifluoroacetic acid, L-cystathionine, PE(18:1), L-arginine, MG(20:4), and dihydroxycholesterol were ranked lower, and linoleic acid, 8-methoxykynurenate, creatine, LysoPC(18:1), L-leucine, L-carnitine, and L-lactic acid were ranked higher in C3H^nDNA^:C57^mtDNA^ MNX compared to C3H^nDNA^:C3H^mtDNA^ WT mice (**Figure 6E, top**). On HFD, 56-dihydroxyprostalglandin-F1a, R-4phosphopantothenoyl-L-cysteine, LysoPC(16:0), 25-dichloro-4-oxohex-2-enedioate, trifluoroacetic acid, L-cystathionine, toluene, MG(20:4), PC(20:1), and PC(16:1) were ranked lower, and linoleic acid, L-histidine, 16-methylheptadecanoic acid, LysoPC(18:1), L-leucine, L-carnitine, arachidonic acid, histamine, creatine, and L-arginine were ranked higher in C3H^nDNA^:C57^mtDNA^ MNX compared to C3H^nDNA^:C3H^mtDNA^ WT mice (**Figure 6E, bottom**).

Overall, the number of differentially abundant plasma metabolites was substantially greater than that of skeletal muscle within each strain. Additionally, in plasma, mice possessing C57^mtDNA^ on the C57BL/6J nuclear background (C57^nDNA^:C57^mtDNA^ WT) displayed > 13 times the number of differentially abundant metabolites in response to HFD than in mice possessing C3H^mtDNA^ on the same nuclear background (C57^nDNA^:C3H^mtDNA^ MNX, 275 vs 21, respectively). Similarly, mice possessing C3H^mtDNA^ on the C3H nuclear background (C3H^nDNA^:C3H^mtDNA^ WT) displayed a ∼92% reduction in the number of total metabolites found in mice possessing C57^mtDNA^ on the same nuclear background (C3H^nDNA^:C57^mtDNA^ MNX, 131 vs 1,627, respectively). Interestingly, this trend was found in untargeted transcriptomic analyses of white adipose tissue (WAT) gene expression, where both epididymal WAT and inguinal WAT stores showed an increased number of differentially expressed genes in response to HFD in mice with C57^mtDNA^ relative to those possessing C3H^mtDNA^ ^52^.

### Metabolite Networks Define Strain-Diet Interactions in CMD

#### Skeletal Muscle

An overview of the Weighted Metabolite Co-Expression Network Analysis derived from the multiWGCNA package is described in **Figure 7A-E**. HFD module names were assigned arbitrary colours (italicized in-text) to differentiate modules. In skeletal muscle, module preservation analysis of mice possessing C57^nDNA^ (C57^nDNA^:C57^mtDNA^ WT and C57^nDNA^:C3H^mtDNA^ MNX) revealed 12 HFD modules not found in control networks (i.e., those with *Z*-Summary < 10) (**Supplementary Table S1A**). Eleven HFD modules were significantly (*P*-adj < 0.05) associated with the C57^nDNA^:C57^mtDNA^ WT strain; of these 11 modules, two were non-preserved and unique to C57^nDNA^:C57^mtDNA^ WT mice (**Supplementary Table S1B, top**). Fourteen HFD modules were significantly (*P*-adj < 0.05) associated with C57^nDNA^:C3H^mtDNA^ MNX mice; of these 14 modules, five were non-preserved and unique only to C57^nDNA^:C3H^mtDNA^ MNX mice (**Supplementary Table S1B, bottom**). HFD-strain module overlaps were calculated, and KEGG-based pathway enrichment analysis was performed on lists of overlapped metabolites.

**Figure 7:**
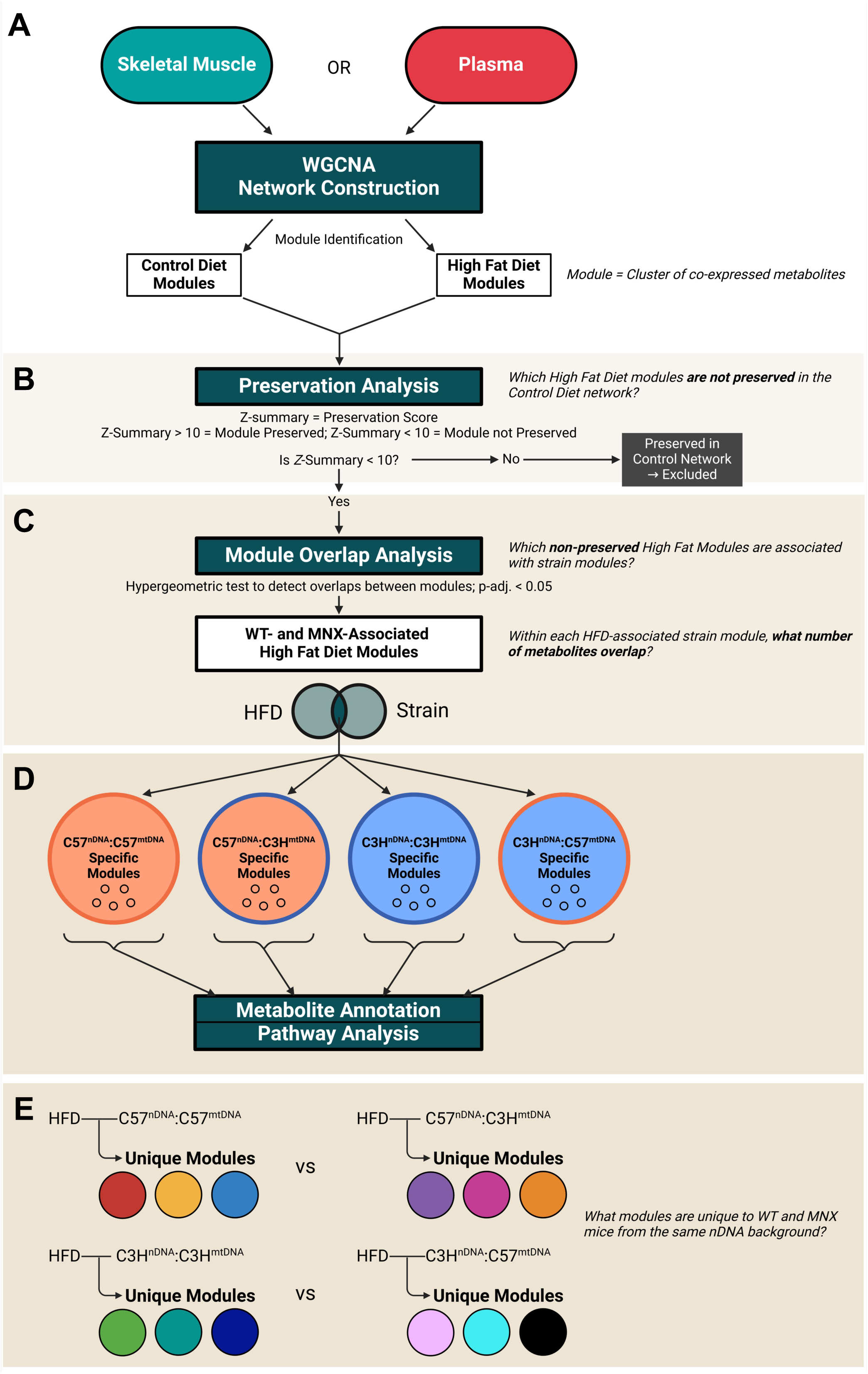
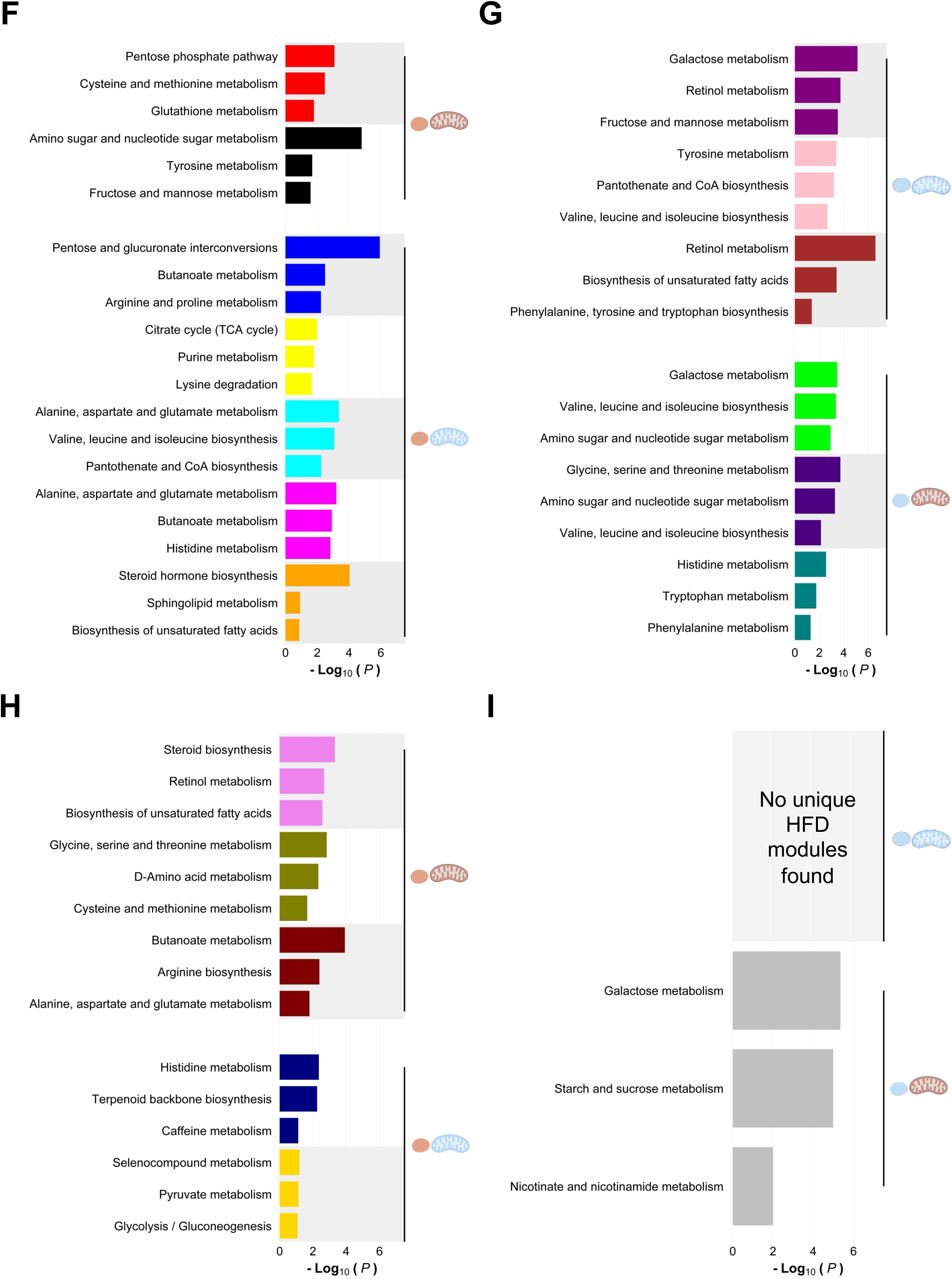
Overview of Weighted Metabolite Co-Expression Network Analysis and Skeletal Muscle and Plasma Pathway Analysis of non-preserved High-Fat Diet (HFD) Modules significantly associated with WT and MNX strains. **(A)** Skeletal Muscle and Plasma networks were independently generated using a novel two-factor strategy, using diet as the primary factor and strain as the secondary factor. Control Diet and HFD Modules were identified. **(B)** Module preservation analysis was performed to evaluate HFD modules that were not found within the Control Diet sub-network. HFD modules with *Z*-Summary values < 10 were deemed non-preserved and are unique to the HFD (disease) state. **(C)** Module overlaps were calculated between strain-specific Wild-Type (WT) and Mitochondrial-Nuclear eXchange (MNX) modules and non-preserved HFD modules. HFD-strain module associations with an adjusted *P*-value < 0.05 were retained. Overlapping metabolites between these significantly associated HFD and strain modules were found. **(D)** HFD-strain overlapping metabolites evaluated from (C) were annotated and contextualized using pathway analysis. **(E)** Unique modules that associated with only WT or only MNX strains were retained and examined. Skeletal Muscle Co-expression Networks outlined: **(F) (top)** two HFD modules (HFD Module Red and Black) significantly associated with C57^nDNA^:C57^mtDNA^ WT mice and **(bottom)** five HFD modules (HFD Module Blue, Yellow, Cyan, Magenta, and Orange) significantly associated with C57^nDNA^:C3H^mtDNA^ MNX mice. **(G) (top)** three HFD modules (HFD Module Purple, Pink, and Brown) significantly associated with C3H^nDNA^:C3H^mtDNA^ WT mice and **(bottom)** three HFD modules (HFD Module Lime, Indigo, and Teal) significantly associated with C3H^nDNA^:C57^mtDNA^ MNX mice. Plasma Co-Expression Networks outlined: **(H) (top)** three HFD modules (HFD Module Violet, Olive, and Maroon) significantly associated with C57^nDNA^:C57^mtDNA^ WT mice and **(bottom)** two HFD modules (HFD Module Navy and Gold) significantly associated with C57^nDNA^:C3H^mtDNA^ MNX mice; **(I) (top)** no HFD modules were significantly associated with C3H^nDNA^:C3H^mtDNA^ WT and **(bottom)** one HFD module (HFD Module Silver) significantly associated with C3H^nDNA^:C57^mtDNA^ MNX mice. Pathway analysis was performed on all modules and the top three pathways by *P*-value are presented. These pathways are representative of metabolite interactions within each module. Modules are coloured arbitrarily to help with identification. Pathway analysis was performed on all modules and the top three pathways by *P*-value are presented. These pathways are representative of metabolite interactions within each module. Modules are coloured arbitrarily to help with identification.

C57^nDNA^:C57^mtDNA^ WT skeletal muscle modules associated with HFD revealed significant enrichment in pathways related to responses to oxidative stress in the *red* module and sugar catabolism in the *black* module (**Figure 7F, top**). In C57^nDNA^:C3H^mtDNA^ MNX mice, pathway enrichment analysis revealed enrichment of carbohydrate, fatty acid, and amino acid metabolism pathways in the *blue* module; citric acid cycle regulation in the *yellow* module; amino acid and vitamin synthesis in the *cyan* module; amino acid and fatty acid metabolism in the *magenta* module; and steroid and sphingolipid metabolism in the *orange* module (**Figure 7F, bottom**).

Skeletal muscle module preservation analysis of mice possessing C3H^nDNA^ (C3H^nDNA^:C3H^mtDNA^ WT and C3H^nDNA^:C57^mtDNA^ MNX) revealed eight HFD modules not found in control networks (**Supplementary Table S2A**). Eleven HFD modules were significantly (*P*-adj < 0.05) associated with the C3H^nDNA^:C3H^mtDNA^ WT strain; of these 11 modules, three were non-preserved and unique to C3H^nDNA^:C3H^mtDNA^ WT mice (**Supplementary Table S2B, top**). Thirteen HFD modules were significantly (*P*-adj < 0.05) associated with the C3H^nDNA^:C57^mtDNA^ MNX strain; of these 13 modules, three were non-preserved and unique to C3H^nDNA^:C57^mtDNA^ MNX mice only (**Supplementary Table S2B, bottom**).

C3H^nDNA^:C3H^mtDNA^ WT skeletal muscle modules associated with HFD revealed significant enrichment in pathways related to retinol and sugar metabolism in the *purple* module; amino acid and vitamin D synthesis in the *pink* module; and fatty acid and aromatic amino acid metabolism in the *brown* module (**Figure 7G, top**). In C3H^nDNA^:C57^mtDNA^ MNX mice, pathway analysis revealed enrichment of sugar and branched chain amino acid metabolism in the *lime* module; polar and branched chain amino acid and sugar metabolism in the Indigo module; and amino acid metabolism in the *teal* module (**Figure 7G, bottom**).

Interestingly, none of the three most enriched pathway terms from the HFD modules in C57^nDNA^:C57^mtDNA^ WT and C57^nDNA^:C3H^mtDNA^ MNX strains were common. This may substantiate the hypothesis that the differing mitochondrial environments of these nDNA-matched strains regulate the response to HFD stress in distinct ways. Only two HFD module-associated pathways were common in C3H^nDNA^-matched strains (C3H^nDNA^:C3H^mtDNA^ WT and C3H^nDNA^:C57^mtDNA^ MNX), including ‘Galactose metabolism’, and ‘Valine, leucine and isoleucine biosynthesis’.

Furthermore, in strains possessing identical C3H^mtDNA^ (C57^nDNA^:C3H^mtDNA^ MNX and C3H^nDNA^:C3H^mtDNA^ WT), three pathways, including ‘Valine, leucine and isoleucine biosynthesis’, ‘Pantothenate and CoA biosynthesis’, and ‘Unsaturated fatty acid biosynthesis’ were common. In C57^mtDNA^-matched mice (C57^nDNA^:C57^mtDNA^ WT and C3H^nDNA^:C57^mtDNA^ MNX), ‘Amino sugar and nucleotide sugar metabolism’ was common.

#### Plasma

In plasma, module preservation analysis of mice possessing C57^nDNA^ (C57^nDNA^:C57^mtDNA^ WT and C57^nDNA^:C3H^mtDNA^ MNX) revealed nine HFD modules not found in control networks (**Supplementary Table S3A**). Fourteen HFD modules were significantly (*P*-adj < 0.05) associated with the C57^nDNA^:C57^mtDNA^ WT strain; of these 14 modules, three were non-preserved and unique to C57^nDNA^:C57^mtDNA^ WT mice (**Supplementary Table S3B, top**). Eleven HFD modules were significantly (*P*-adj < 0.05) associated with the C57^nDNA^:C3H^mtDNA^ MNX strain; of these 11 modules, two were non-preserved and unique to C57^nDNA^:C3H^mtDNA^ MNX mice (**Supplementary Table S3B, bottom**).

C57^nDNA^:C57^mtDNA^ WT plasma modules associated with HFD revealed significant enrichment in pathways broadly related to the steroid and fatty acid biosynthesis in the *violet* module; polar and sulphur-containing amino acid metabolism in the *olive* module; and butanoate and amino acid metabolism in the *maroon* module (**Figure 7H, top**). In C57^nDNA^:C3H^mtDNA^ MNX mice, pathway analysis revealed enrichment of the metabolism of histidine and terpenoid metabolism pathways in the *navy* module and glucose oxidation pathways in the *gold* module (**Figure 7H, bottom**).

Plasma module preservation analysis of mice possessing C3H^nDNA^ (C3H^nDNA^:C3H^mtDNA^ WT and C3H^nDNA^:C57^mtDNA^ MNX) revealed three HFD modules not found in control networks (**Supplementary Table S4A**). Four HFD modules were significantly (*P*-adj < 0.05) associated with the C3H^nDNA^:C3H^mtDNA^ WT strain; unlike other network analyses, of these four modules, none were non-preserved and unique to C3H^nDNA^:C3H^mtDNA^ WT mice (**Supplementary Table S4B, top; Figure 7I, top**). Seven HFD modules were significantly (*P*-adj < 0.05) associated with the C3H^nDNA^:C57^mtDNA^ MNX strain; of these seven modules, one was non-preserved and unique to C3H^nDNA^:C57^mtDNA^ MNX mice only (**Supplementary Table S4B, bottom**). Pathway enrichment analysis of this module revealed sugar metabolism pathways in the *gray* module (**Figure 7I, bottom**). In contrast to skeletal muscle, there were no common pathways in nDNA-matched and mtDNA-matched strains.

## Discussion

The management of CMD remains a challenge due to the interactions between environmental, genetic, and lifestyle factors that modulate CMD susceptibility ^7^. Ectopic fat deposition during HFD feeding accelerates the development of systemic and skeletal muscle insulin resistance; this relationship involves dysregulated mitochondrial fatty acid oxidation ^107–111^. Chronic HFD consumption leads to metabolic inflexibility: the reduced ability to switch from fatty acid to glucose oxidation during increased nutritional intake. Mitochondria are hubs of metabolism within the cell. Point mutations and SNPs in mtDNA are implicated in the pathogenesis of mitochondrial diseases, including diabetic and cardiovascular disease phenotypes ^49,53,54,112,113^. Therefore, it has been proposed that the individual susceptibility to these diseases in conditions of HFD consumption is modulated by mtDNA background ^31,52,75,114–116^. Models isolating the effect of mtDNA and nDNA have demonstrated that distinct mtDNA sequences can alter baseline insulin sensitivity and glucose tolerance in cells and muscle tissue ^47,112,117–119^. However, the metabolites that drive these mtDNA-based differences are largely unknown.

We investigated metabolites and metabolite networks that may be differentially regulated through interactions between mtDNA and diet. Using the MNX model, we evaluated skeletal muscle- and plasma-specific metabolomic profiles of WT and MNX mice during HFD feeding. A summary of these results is presented in the graphical abstract (**Figure 8)**. Previous studies of diet-induced CMD in the MNX mouse model have demonstrated a differential susceptibility to CMD development that more closely aligns with their mtDNA background ^47,52,73,75,120^. Specifically, C57^mtDNA^ induces a more pronounced obesity phenotype, pathogenic WAT gene expression profiles, elevates hepatic liver fibrosis, and reduces systemic and skeletal muscle insulin sensitivity.

**Figure 8:**
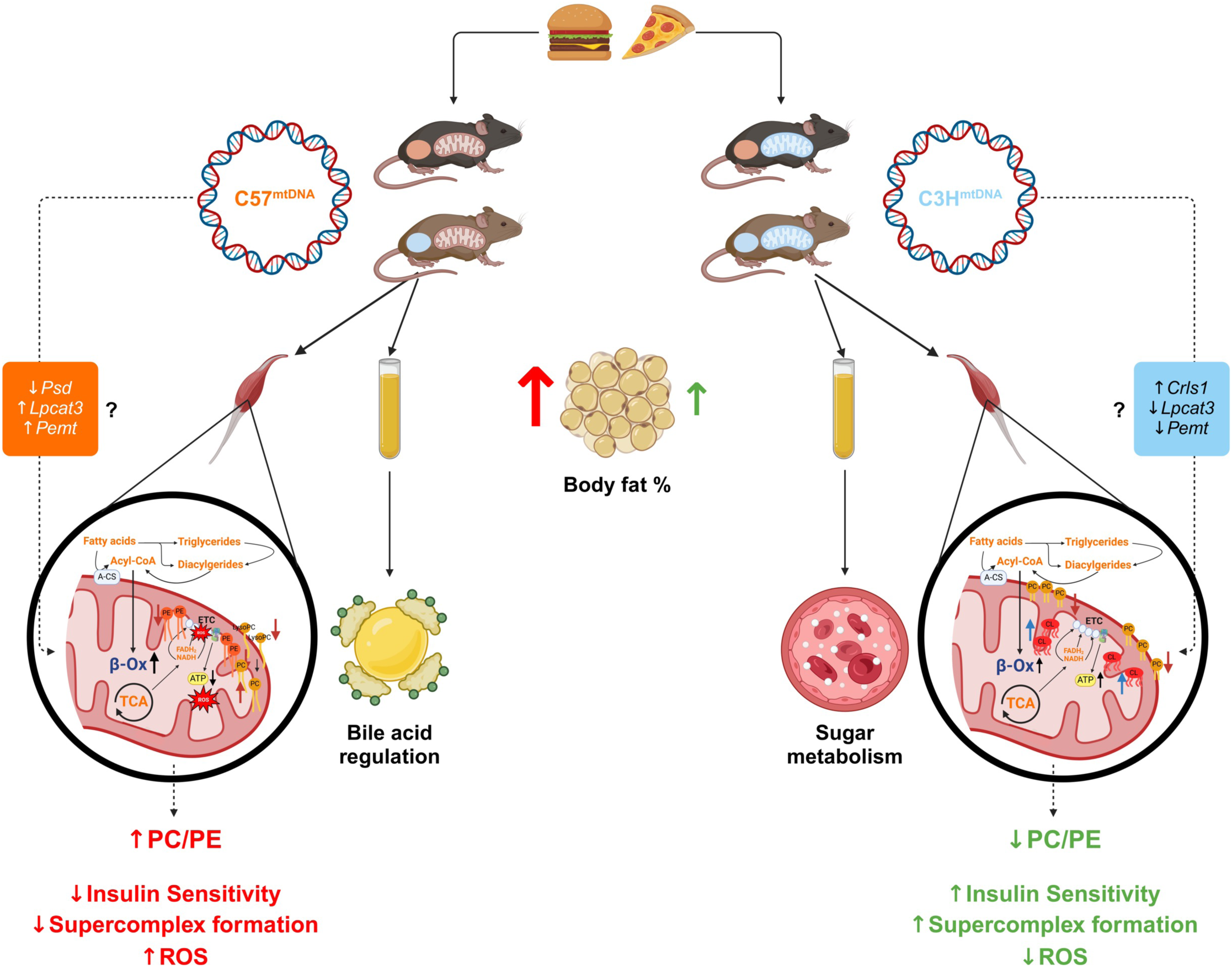
Summary of results from multi-tissue metabolomics analysis in the MNX mouse model. When fed the same high-fat diet, mice possessing C57^mtDNA^ demonstrated higher body fat % relative to those mice possessing C3H^mtDNA^. Within skeletal muscle, reduced levels of glycerophospholipids including phosphatidylethanolamines (PEs) and lysophosphatidylcholines (LysoPCs) and elevated levels of phosphatidylcholines (PCs) were observed in mice possessing C57^mtDNA^; in mice possessing C3H^mtDNA^, cardiolipin (CLs) were elevated and PCs were reduced in abundance. These results point towards an elevated PC/PE ratio in cardiometabolic disease (CMD)-susceptible mice possessing C57^mtDNA^ and a depressed ratio in CMD-resistant mice possessing C3H^mtDNA^, which may explain previously reported differences in skeletal muscle insulin sensitivity and cellular ROS production between mice with distinct mtDNA backgrounds. Several nuclear-encoded genes including phosphatidylserine decarboxylase (*Psd*), lysophosphatidylcholine acyltransferase 3 (*Lpcat3*), phosphatidylethanolamine N-methyltransferase (*Pemt*), and cardiolipin synthase 1 (*Crls1*) may drive changes in abundance to these glycerophospholipids. In plasma, significant changes to bile acid regulation and biosynthesis and sugar metabolism-related pathways were found in mice possessing C57^mtDNA^ and C3H^mtDNA^, respectively. Gut-liver-muscle signaling through modulation of bile acids may induce insulin resistance phenotypes in CMD-susceptible mice, whereas this was not observed in CMD-resistant mice.

Glycerophospholipids, specifically PCs and PEs, were consistently differentially abundant in both skeletal muscle and plasma. Additionally, PCs and glycerophosphocholines (precursors to PCs and PEs) had distinct rank changes at baseline in all mice. PCs and PEs are membrane phospholipids abundant within the plasma membrane and both mitochondrial outer and inner membranes ^121^. Unlike PCs, PEs do not participate in bilayer formation and function to stabilize the mitochondrial membranes ^121^. In the skeletal muscle of mice with C57^nDNA^, PCs were generally downregulated in mice possessing C3H^mtDNA^, whereas PEs were downregulated in mice possessing C57^mtDNA^. The ratio of PC/PE has emerged as a validated indicator of muscle insulin sensitivity and functional activity in exercise and diabetes models ^122–126^. Although increases in muscular PC and PE levels are indicators of preserved insulin signalling, the PC/PE ratio is negatively correlated with insulin sensitivity ^126^. Our results suggest that the PC/PE ratio is raised in mice harboring C57^mtDNA^ (C57^nDNA^:C57^mtDNA^ WT and C3H^nDNA^:C57^mtDNA^ MNX) and lowered in mice harboring C3H^mtDNA^ (C3H^nDNA^:C3H^mtDNA^ WT and C57^nDNA^:C3H^mtDNA^ MNX). Sarcoplasmic reticulum Ca^2+^-ATPase (SERCA) function, which regulates skeletal muscle relaxation and muscle mass, is inversely correlated with the PC/PE ratio ^125,127–129^. SERCA activity also plays a role in maintaining lipogenesis in muscle, which may influence the propensity to develop lipid-induced insulin resistance ^130^. Additionally, mitochondrial supercomplex assembly of Complexes I and IV is impaired and mitochondrial fragmentation is increased in PE-deficient mitochondria ^131,132^. Mice with PE deficiency develop obesity and insulin resistance leading to impaired glucose uptake ^100^. This is concordant with decreased glucose levels in the muscle of C57^nDNA^:C57^mtDNA^ WT mice. Although a bioenergetic analysis of skeletal muscle in the MNX mouse model has not yet been published, reports in cardiac muscle show decreased transcription of genes involved in bioenergetic pathways in mice with C57^mtDNA^ in response to volume overload ^52,120^. Therefore, the rise in PC/PE may induce similar bioenergetic disruptions, potentially mediated by oxidative stress as seen in previous reports ^73,75,120,133^. The activity of phosphatidylserine decarboxylase (Psd), the enzyme which synthesizes PE, and phosphatidylethanolamine *N*-methyltransferase (Pemt) which catalyzes the conversion of PE to PC, also may be modulated by differences in mtDNA background; Psd deficiency models have displayed substantial increases in oxidative stress ^134^. Therefore, further investigations of the mitochondria from C57BL/6J mice should investigate the activity of Psd relative to the mitochondria of C3H/HeN mice. Bioenergetic disruptions are a known feature of insulin resistance ^135–137^, and therefore, modulation of the PC/PE ratio presents a potential mechanism by which mtDNA primes the activity of oxidative stress-inducing enzymes such as Psd.

Further, LysoPC, a metabolic precursor to PC, was downregulated in the skeletal muscle of mice harboring C57^mtDNA^. LysoPC is vital in plasma membrane organization, and overexpression of the LysoPC-to-PC conversion enzyme LysoPC acyltransferase 3 (Lpcat3) has been demonstrated to promote glucose intolerance and insulin resistance in myocytes ^138^. Additionally, the generation of muscle force and the size of muscle fibers is reduced in mice with low skeletal muscle LysoPC, which is exacerbated by HFD feeding ^139^. Although PCs were not found to be increased in the skeletal muscle of C57^mtDNA^ mice concordant with a decrease in LysoPC, lowering LysoPC levels alone can predict significant changes in tissue-level insulin resistance ^138^. Additionally, in the liver, reduced LysoPC levels may be indicative of increased fatty acid oxidation; therefore, muscle LysoPC levels may indicate enhanced beta-oxidation as a mechanism of lipid-induced insulin resistance ^140,141^.

We have also found that CLs were upregulated in the skeletal muscle of mice harboring C3H^mtDNA^, however a clear regulation pattern based on genetic background could not be ascertained in plasma. CL is a phospholipid primarily found in the inner mitochondrial membrane and has a vital role in maintaining the organization of mitochondrial supercomplexes for efficient mitochondrial coupling ^142^. Increased CL levels are indicative of the maintenance of ATP coupling rates in the face of HFD-induced oxidative stress in C3H^mtDNA^ ^143^. CL has been demonstrated to maintain lipid oxidative capacity through reductions in overall ATP production; this may be a sign of lower mitochondrial economy as a protective measure to reduce total ROS levels in mice harboring C3H^mtDNA 115,143^.

Interestingly, alanine was consistently downregulated in the skeletal muscle of mice harboring C3H^mtDNA^, but not clearly differentially regulated in mice harboring C57^mtDNA^. Alanine is a non-essential amino acid that participates in the regulation of gluconeogenesis and amino acid turnover, through recycling by the liver and muscle ^144^. During fasting, alanine is transported to the liver through alanine aminotransferase, mediated by peroxisome proliferator-activated receptor gamma coactivator 1-alpha ^145^. Disrupted alanine and glucose transport between the liver and skeletal muscle has been implicated in skeletal muscle CMD phenotypes in *ob*/*ob* mice ^146,147^. Although the regulation of alanine in this study does not neatly align with a particular mtDNA background, its depletion in C3H^mtDNA^ may implicate reduced liver-muscle crosstalk in modulating gluconeogenesis in times of HFD feeding. Furthermore, contrary to previous reports of DG accumulation in HFD stress ^148,149^, DGs were downregulated in HFD feeding in both C57^nDNA^:C57^mtDNA^ WT and C57^nDNA^:C3H^mtDNA^ MNX skeletal muscle. This may indicate adequate catabolism of DGs into free fatty acids, but impaired beta-oxidation as proposed as a model of lipid-induced insulin resistance. Also, DGs are involved in the generation of PCs and PEs as a choline recipient, catalyzed by the enzyme CDP-choline:1,2-diacylglycerol choline/ethanolamine phosphotransferase, and therefore reductions in DG levels may influence the ratio of PCs and PEs in muscle ^132,150,151^.

The Weighted Metabolite Co-Expression Network Analysis of skeletal muscle demonstrated that nDNA-mtDNA interactions produce unique metabolite modules associated with diverse metabolic activities. In C57^nDNA^:C57^mtDNA^ WT skeletal muscle, modules related to oxidative stress responses and sugar metabolism were found. Previous studies have demonstrated that mice with C57^mtDNA^ have increased ROS production at baseline, and therefore the co-expression of metabolites involved in glutathione and pentose phosphate pathway may act as a regulatory response to reduce excessive oxidative stress ^73,75,120^. Interestingly, we saw none of the same pathways in C57^nDNA^:C3H^mtDNA^ MNX muscle, which exhibited a diverse set of pathways perhaps involved in the stable regulation of metabolites involving the citric acid cycle including pyruvate metabolism which is central to the regulation of glycolysis and amino acid interconversion ^152^. In C3H^nDNA^:C3H^mtDNA^ WT muscle, the network analysis demonstrated differential amino acid metabolism in branched-chain amino acids and fatty acids, which shows conserved modules in mice with the same C3H^mtDNA^. In C3H^nDNA^:C57^mtDNA^ MNX mice, the network analysis demonstrated similar sugar metabolism-related pathways as seen in C57^nDNA^:C57^mtDNA^ WT mice, again displaying mtDNA-dependent co-regulation of sugar metabolism. We propose that these findings may be related to alterations in insulin signalling inhibiting glucose oxidation in the CMD state, previously displayed in MNX mice ^47^. In the plasma of these mice, we saw more widely varying profiles, with no similar patterns of module expression in WT and MNX strains. This may imply that unique nDNA-mtDNA interactions pose unique plasma co-expression profiles that cannot be aligned with nDNA or mtDNA backgrounds alone.

In plasma, although PCs and PEs were differentially abundant, the absolute ratio of PC/PE in the current study is unable to be determined. However, it has recently been observed that patients with cardiovascular disease, who also present with an increased risk of insulin resistance development, have *lower* circulating PC/PE ratios compared to controls. The plasma PC/PE ratio is also inversely associated with nucleotide-binding oligomerization domain-like receptor family pyrin domain-containing 3 (NLRP3) levels ^153^. Additionally, cardiac left ventricle remodelling was correlated with the increased levels of NLRP3 in these subjects ^153^. As systemic inflammation and left ventricular damage have been observed in mice harboring C57^mtDNA^ ^73,75,120^, it is important for future studies to gauge circulating PC and PE levels to gauge their contribution to other multi-organ CMD pathologies. Additionally, plasma profiles in mice harboring C57^mtDNA^ exhibited downregulation of various CDP-DGs. In the liver, mammalian target of rapamycin complex 1 regulation controls the secretion of phospholipids and fatty acids incorporated in very low-density lipoproteins ^154^. CDP-DG is a crucial mediator of obesity development that is controlled by transcriptional regulation in the liver, so further studies of the liver and skeletal muscle axis in MNX mice must be conducted to understand how circulating CDP-DG levels are reflected in the skeletal muscle ^155^.

In general, we saw variation in the glycerophospholipid synthesis pathways in plasma of all strains, especially involving CDP-DGs, PAs, CLs, and phosphoinositols. Future studies using a targeted lipidomics approach in the plasma of WT and MNX mice may glean a clearer picture of changes in particular subgroups of these phospholipids that align with CMD signature. Due to the abundance of differentially regulated metabolites in plasma relative to skeletal muscle, pathway analysis was performed and revealed perturbations to primary bile acid metabolism that were found in mice harboring C57^mtDNA^ (C57^nDNA^:C57^mtDNA^ WT and C3H^nDNA^:C57^mtDNA^ MNX) and sugar and glycolysis related pathways in mice with C3H^mtDNA^ (C3H^nDNA^:C3H^mtDNA^ WT and C57^nDNA^:C3H^mtDNA^ MNX). Although the systemic regulation of bile acids has not been deduced in studies of MNX mice, perturbed bile acid production has been noted in CMD development through alterations in immunometabolism ^156^. Bile acids and associated metabolites are modified by gut microbiota and are known to interact with the Farnesoid X Receptor to induce sarcopenic and insulin-resistant phenotypes in muscle ^157–159^. Therefore, bile acid dysregulation and potential dysbiosis in the gut microbiota, or alterations in liver secretion of bile acids, play a role in modulating systemic CMD profiles. Interestingly, liver transcriptomics in MNX mice show differences in the regulation of genes related to inflammation, and fibrosis; the altered expression of lipid metabolism mediators such as peroxisome proliferator-activated receptor-gamma during HFD consumption is known to dysregulate hepatic bile acid production ^73,160^. It is known that bile acids may indirectly modulate skeletal muscle fatty acid oxidation and volume, which regulate insulin signalling ^42,161–163^. As insulin signalling profiles of mice harboring C3H^mtDNA^ are known to reflect lower levels of plasma glucose and insulin, as well as healthy phosphorylated-protein kinase B to total protein kinase B ratios in skeletal muscle ^47^, the lowered abundance of glucose and carbohydrate metabolites in plasma may reflect healthy import into highly glycolytic tissues.

The plasma Weighted Metabolite Co-Expression Network Analysis revealed only one unique module in C3H^nDNA^:C57^mtDNA^ MNX mice and no unique modules in C3H^nDNA^:C3H^mtDNA^ WT mice, perhaps displaying a lack of metabolite co-regulation in plasma when mice with C3H^nDNA^ are faced with HFD. The reason for this is not known. In mice possessing C57^nDNA^ (C57^nDNA^:C57^mtDNA^ WT and C57^nDNA^:C3H^mtDNA^ MNX), the network analysis revealed changes in steroid, amino acid, and butanoate metabolism in C57^nDNA^:C57^mtDNA^ WT mice and terpenoid biosynthesis and glycolytic metabolism in C57^nDNA^:C3H^mtDNA^ MNX mice. An enhanced circulation of steroid hormones is implicated in CMD-associated cardiovascular diseases such as hypertension ^164^. It seems here that differences in mtDNA and potential nDNA-mtDNA interactions coordinate co-regulation profiles with the same C57^mtDNA^. However, the cause of the relative lack of metabolite co-regulation in mice possessing C3H^nDNA^ relative to C57^nDNA^ should be further investigated.

## Conclusions

Overall, this study revealed unique metabolomic profiles in the skeletal muscle and plasma of nDNA-matched WT and MNX mice. CMD is a complex array of risk factors involving the coordination of multiple tissues including the liver, pancreas, gut, brain, adipose, endocrine, and skeletal muscle organs. These results suggest that glycerophospholipid profiles in skeletal muscle promotes electron transport chain dysfunction through a profile favouring abnormally increased beta-oxidation in C57^mtDNA^ mice and relative protection against elevated beta-oxidation in mice possessing C3H^mtDNA^. In the plasma, bile acid and sugar metabolism pathways are differentially regulated in mice possessing C57^mtDNA^ and C3H^mtDNA^, respectively.

Our work suggests that mtDNA is a mediating factor in the differential development of CMD, complementary to nDNA and diet. Although the nDNA accounts for 99% of the proteins found in the mitochondria ^165^, it has been proposed that the expression of these nuclear genes is at least in part modulated by mtDNA signature through changes in metabolites. Our studies propose that C57^mtDNA^ may ‘prime’ the mitochondrial environment to show disease-susceptible profiles in skeletal muscle and plasma in response to HFD. As skeletal muscle is often the first tissue to undergo insulin resistance in the formation of CMD, it presents an attractive marker to predict future susceptibility to disease based on dietary, genetic, and environmental factors ^166,167^. Further investigations into the metabolites outlined here and the flux of metabolites regulating transcriptional changes in skeletal muscle will provide us with greater insight into nDNA-mtDNA interactions and avenues to treat multi-organ CMD before progression into cardiovascular disease and T2DM.

## Limitations and Future Work

While the modulation of glycerophospholipid abundance appears to be the predominant change in the skeletal muscle metabolome in our mouse model, we did not quantify total PC and PE levels through the untargeted metabolomics approach; additionally, we did not assess the abundance of glycerophospholipids with varying chain lengths. Future investigations should address these parameters to clarify their potential roles in modulating beta-oxidation rates. Additionally, the relatively lower number of metabolites detected in skeletal muscle compared to plasma may reflect differences in metabolome complexity, though sample degradation during collection or preprocessing could also have contributed to this disparity.

Furthermore, this study is limited to male mice fed a low-sucrose, HFD over a relatively short duration. Constraints in breeding colony maintenance and the high cost of analyses precluded the inclusion of female mice. Expanding future research to include female mice is critical for drawing more comprehensive and generalizable conclusions. Similarly, extending the duration of diet exposure and systematically varying diet composition will provide deeper insights into how these factors influence metabolomic profiles. Investigating these variables in greater detail will help delineate the specific effects of diet composition and duration on skeletal muscle metabolism.

## Acknowledgements

The authors wish to thank the Clinical Biomarkers Laboratory staff at Emory University for preprocessing samples, extracting HPLC-MS data, and providing analytical support. We also thank the Translational Institute of Medicine (TIME) and the staff at the Queen’s CardioPulmonary Unit (QCPU) for their advice and input.

## Author contributions

Conceptualization: K.DS.

Methodology: A.S., K.DS., C.C.T.H., M.R.S.

Investigation: A.S., M.S.W., D.M.M., M.K., J.L.M.H.V., M.R.S.

Visualization: A.S., K.DS., C.C.T.H.

Formal Analysis: A.S., M.R.S.

Funding acquisition: K.DS.

Project administration: A.S., K.DS., M.R.S.

Resources: K.DS., C.C.T.H., M.R.S.

Software: A.S., C.C.T.H., M.R.S.

Supervision: K.DS., C.C.T.H.

Writing – original draft: A.S., K.DS., C.C.T.H.

Writing – review & editing: A.S., K.DS., C.C.T.H.

## Data and materials availability

All data and scripts needed to perform the analyses in this paper are available upon reasonable request from the corresponding author.

## Supplementary Materials

Figs. S1 to S2

Tables S1 to S4

---

**Figure S1:**
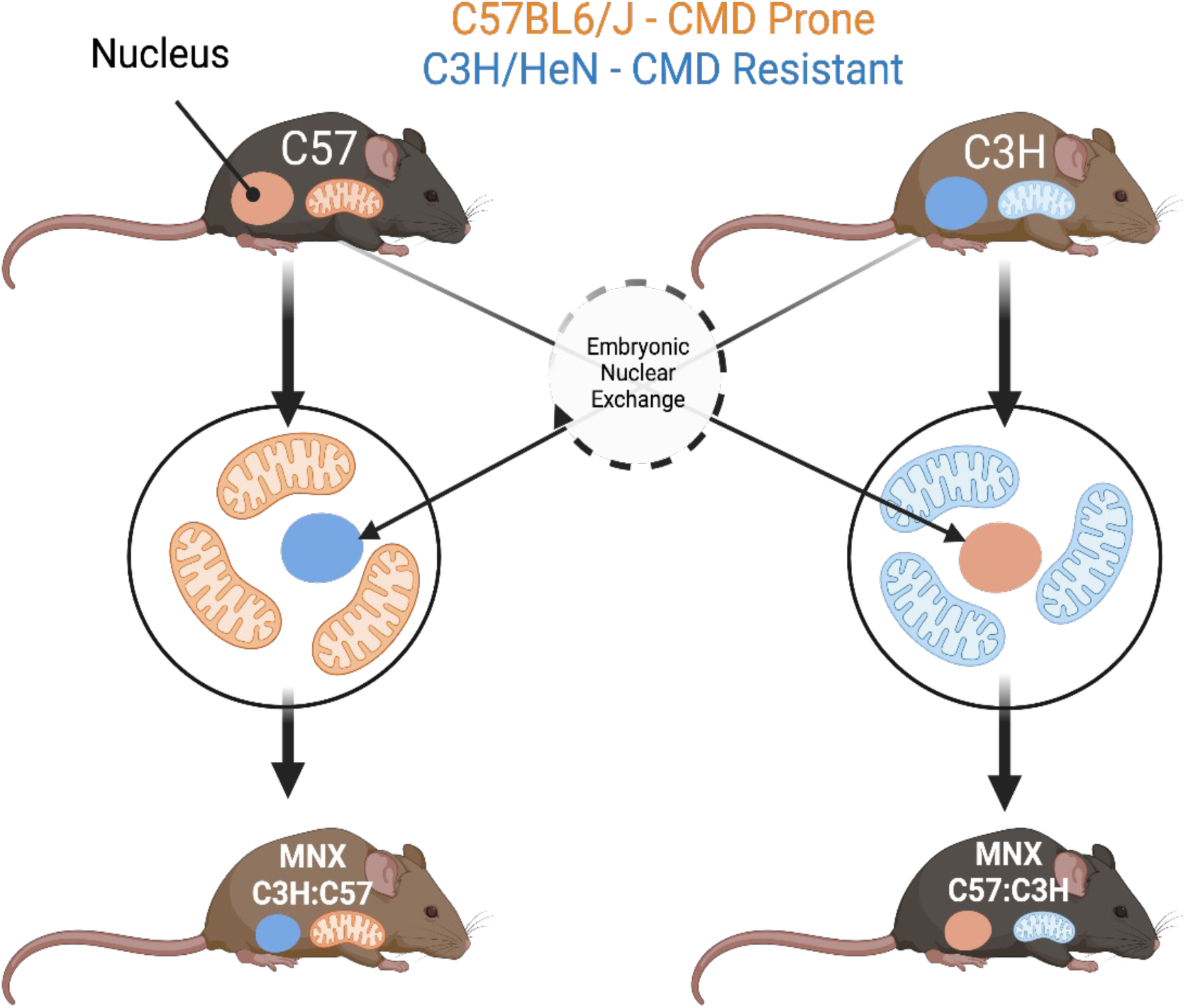
Overview of the Mitochondrial-Nuclear eXchange (MNX) mouse model. C57BL/6J (C57) and C3H/HeN (C3H) inbred mouse strains possess known differential susceptibilities to cardiometabolic disease (CMD) development. C57 mice are CMD-prone while C3H mice are CMD-resistant. To assess the contribution of mitochondrial DNA to CMD pathogenesis, the pro-nuclei of C57 and C3H embryos are reciprocally exchanged. This exchange results in MNX mice with mismatched nuclear and mitochondrial genetic backgrounds.

**Figure S2:**
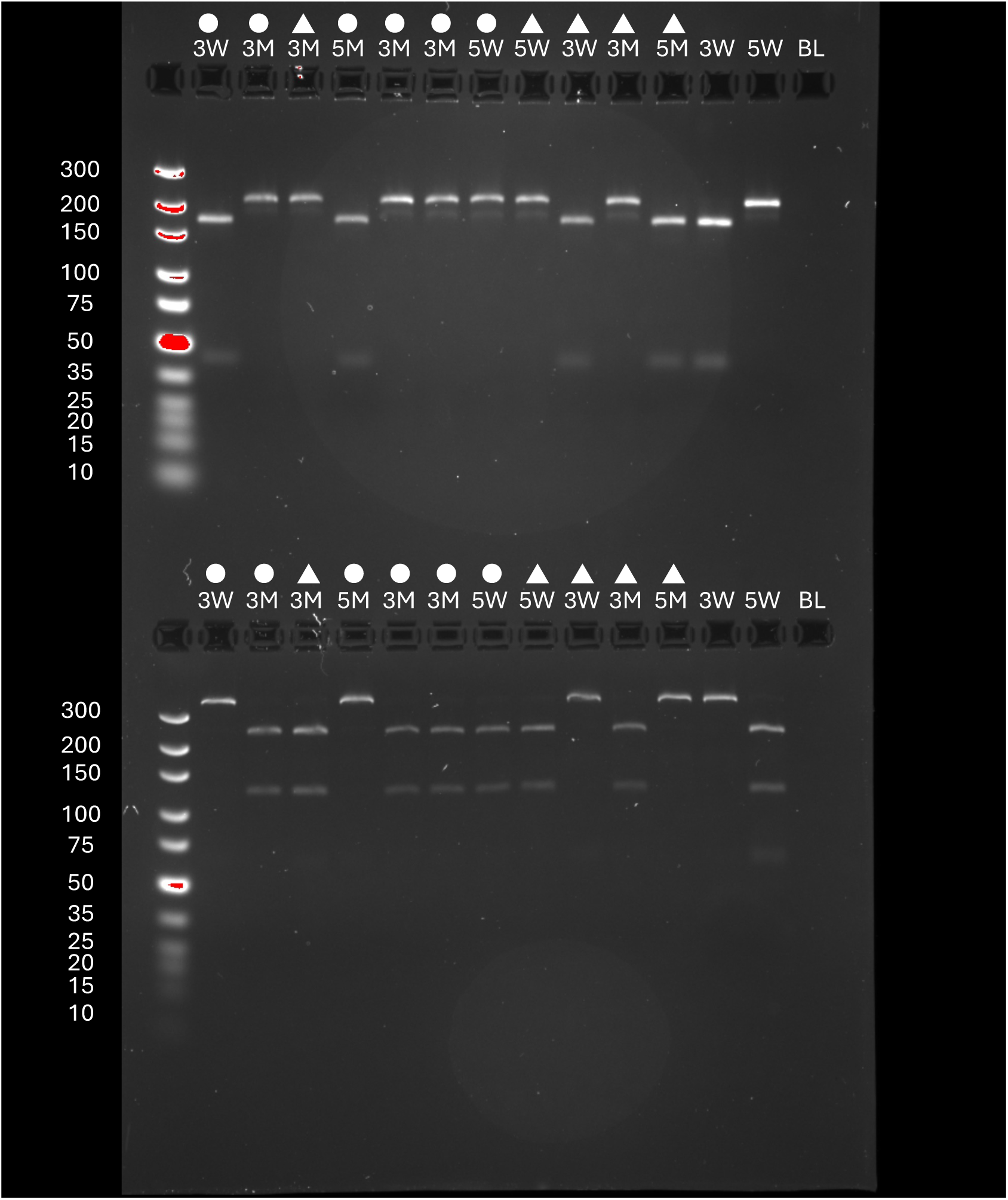
Representative haplotyping gel for validation of pro-nuclear transfer, in generation of Mitochondrial-Nuclear eXchange (MNX) mice. (Top) C3H^nDNA^:C3H^mtDNA^ WT (‘3W’) and C57^nDNA^:C3H^mtDNA^ MNX (‘5M’) mice possess the same C3H^mtDNA^. *Bcl1* cleaves C3H^mtDNA^ at bp 9461 into 166 bp and 38 bp fragments but leaves C57^mtDNA^ intact (found in C57^nDNA^:C57^mtDNA^ WT (‘5W’) and C3H^nDNA^:C57^mtDNA^ MNX (‘3M’) mice). Circles represent Control Diet samples and triangles represent High-Fat Diet (HFD) samples. 5W, 3W, and blank (BL) negative control wells are included. (Bottom) C57^nDNA^:C57^mtDNA^ (‘5W’) and C3H^nDNA^:C57^mtDNA^ (‘3M’) mice possess the same C57^mtDNA^. *Pflf1* cuts C57^mtDNA^ at bp 9348 into 274 bp and 111 bp fragments but leaves C3H^mtDNA^ intact (found in C3H^nDNA^:C3H^mtDNA^ (‘5W’) and C57^nDNA^:C3H^mtDNA^ (‘5M’) mice). Circles represent Control Diet samples and triangles represent HFD samples. 5W, 3W, and blank (BL) negative control wells are included.

**Table S1:** Results of overlap analysis and module preservation analysis within the Skeletal Muscle HFD network in mice possessing C57^nDNA^. (A) Results of module preservation analysis of High-Fat Diet (HFD) modules not found in Control Diet network. *Z*-Summary values are ranked and modules with values < 10 were retained. Module numbers are arbitrary labels. (B) HFD modules significantly associated with C57^nDNA^:C57^mtDNA^ WT (‘C57WT’) and C57^nDNA^:C3H^mtDNA^ MNX (‘C57MNX’) strains. Only HFD modules that are unique to the WT or MNX strain were investigated, not those that are shared by both strains. Adjusted *P*-value represents significance of association. Coloured modules are those represented in Figure 7.

| HFD | Module Size | Z-Summary Value |
| --- | --- | --- |
| HFD_012 | 74 | 0.635 |
| HFD_019 | 47 | 1.507 |
| HFD_000 | 647 | 1.596 |
| HFD_011 | 75 | 3.324 |
| HFD_016 | 56 | 3.771 |
| HFD_017 | 56 | 4.593 |
| HFD_006 | 110 | 4.614 |
| HFD_010 | 78 | 4.875 |
| HFD_020 | 46 | 6.416 |
| HFD_009 | 84 | 6.879 |
| HFD_018 | 54 | 7.37 |
| HFD_015 | 63 | 7.615 |
| HFD_004 | 133 | 9.872 |
| HFD_002 | 146 | 12.35 |
| HFD_014 | 70 | 12.41 |
| HFD_007 | 105 | 13.02 |
| HFD_005 | 111 | 13.15 |
| HFD_008 | 98 | 14.93 |
| HFD_003 | 143 | 16.66 |
| HFD_013 | 74 | 18.97 |
| HFD_001 | 229 | 27.5 |

B
| HFD | C57WT | Adjusted P-Value |
| --- | --- | --- |
| HFD_001 | C57WT_002 | 4.73E-73 |
| HFD_007 | C57WT_005 | 2.87E-65 |
| HFD_011 | C57WT_016 | 1.55E-45 |
| HFD_010 | C57WT_009 | 5.15E-39 |
| HFD_000 | C57WT_000 | 1.30E-37 |
| HFD_009 | C57WT_007 | 1.29E-28 |
| HFD_019 | C57WT_008 | 2.08E-24 |
| HFD_013 | C57WT_003 | 4.11E-22 |
| HFD_002 | C57WT_001 | 4.95E-19 |
| HFD_014 | C57WT_004 | 5.49E-07 |
| HFD_012 | C57WT_012 | 0.001751 |

| HFD | C57MNX | Adjusted P-Value |
| --- | --- | --- |
| HFD_001 | C57MNX_001 | 1.01E-65 |
| HFD_004 | C57MNX_002 | 3.80E-37 |
| HFD_006 | C57MNX_017 | 1.46E-35 |
| HFD_000 | C57MNX_000 | 4.74E-31 |
| HFD_007 | C57MNX_012 | 4.15E-23 |
| HFD_014 | C57MNX_006 | 1.19E-22 |
| HFD_002 | C57MNX_003 | 1.24E-20 |
| HFD_003 | C57MNX_004 | 1.10E-17 |
| HFD_020 | C57MNX_009 | 2.44E-14 |
| HFD_015 | C57MNX_013 | 5.49E-11 |
| HFD_012 | C57MNX_018 | 1.01E-08 |
| HFD_009 | C57MNX_008 | 2.37E-07 |
| HFD_018 | C57MNX_007 | 1.80E-05 |
| HFD_019 | C57MNX_016 | 0.000767 |

**Table S2:**
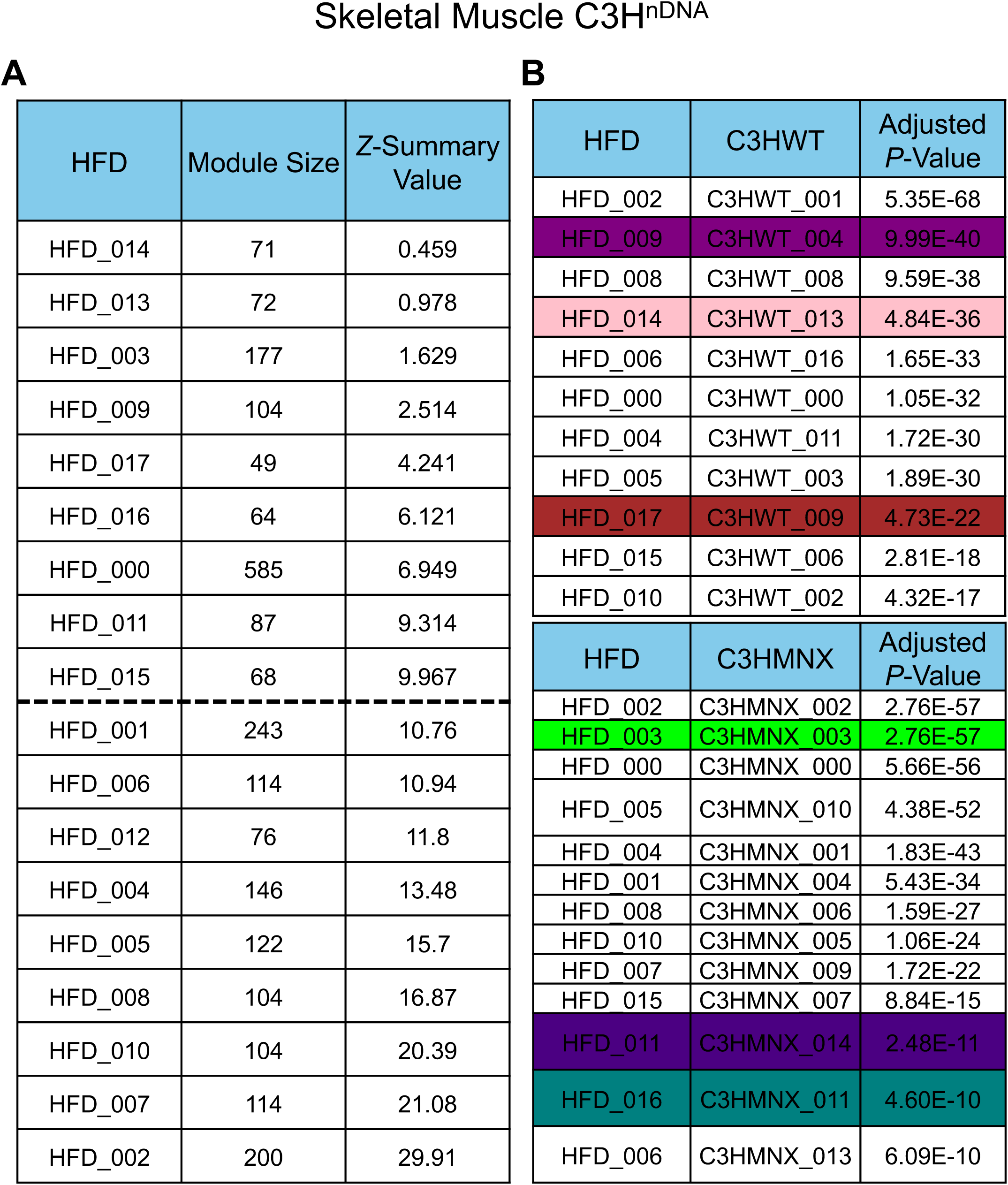
Results of overlap analysis and module preservation analysis within the Skeletal Muscle HFD network in mice possessing C3H^nDNA^. (A) Results of module preservation analysis of High-Fat Diet (HFD) modules not found in Control Diet network. *Z*-Summary values are ranked and modules with values < 10 were retained. Module numbers are arbitrary labels. **(B)** HFD modules significantly associated with C3H^nDNA^:C3H^mtDNA^ WT (‘C3HWT’) and C3H^nDNA^:C57^mtDNA^ MNX (‘C3HMNX’) strains. Only HFD modules that are unique to the WT or MNX strain were investigated, not those that are shared by both strains. Adjusted *P*-value represents significance of association. Coloured modules are those represented in Figure 7.

**Table S3:**
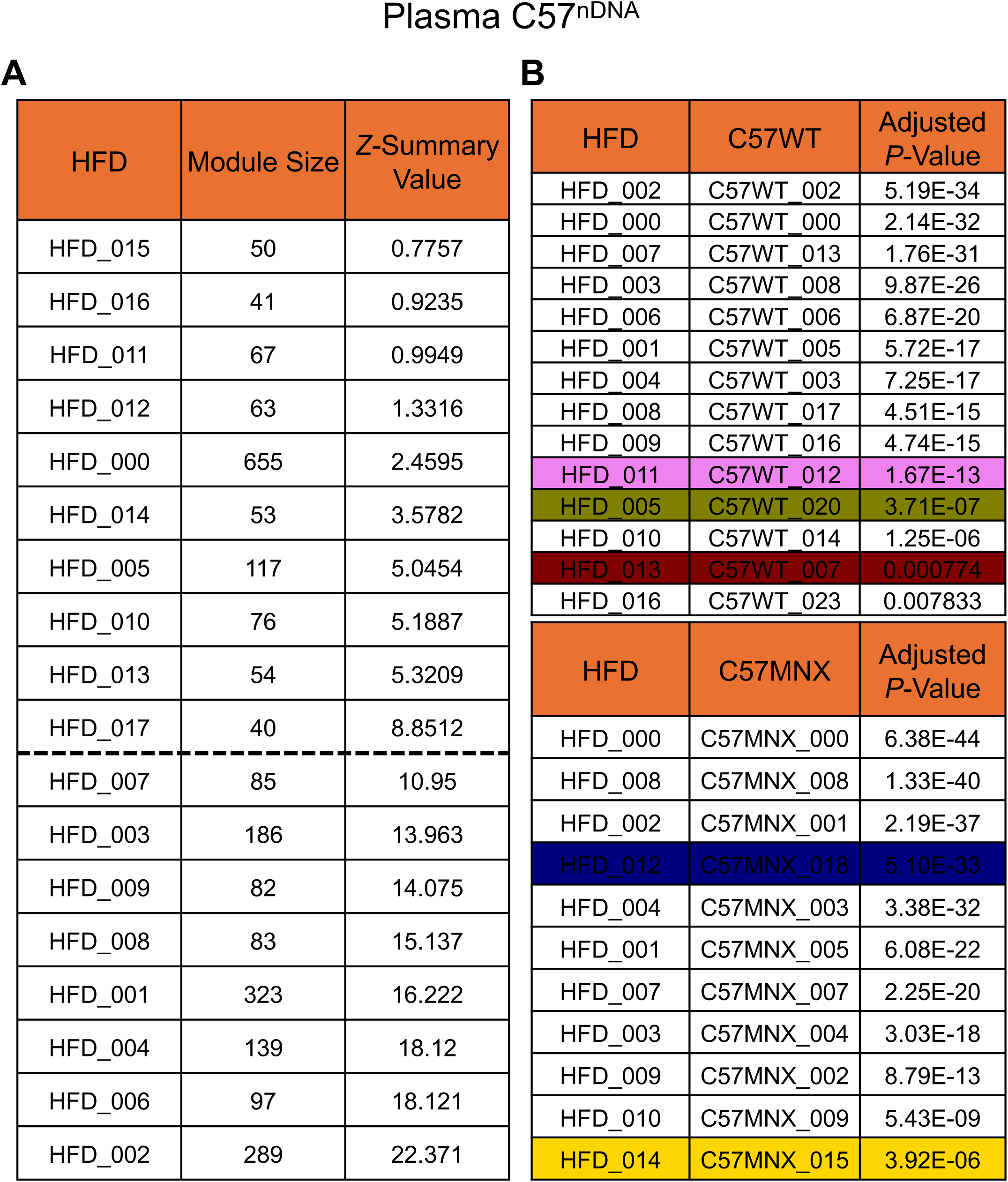
Results of overlap analysis and module preservation analysis within the Plasma HFD network in mice possessing C57^nDNA^. (A) Results of module preservation analysis of High-Fat Diet (HFD) modules not found in Control Diet network. *Z*-Summary values are ranked and modules with values < 10 were retained. Module numbers are arbitrary labels. **(B)** HFD modules significantly associated with C57^nDNA^:C57^mtDNA^ WT (‘C57WT’) and C57^nDNA^:C3H^mtDNA^ MNX (‘C57MNX’) strains. Only HFD modules that are unique to the WT or MNX strain were investigated, not those that are shared by both strains. Adjusted *P*-value represents significance of association. Coloured modules are those represented in Figure 7.

**A**
| HFD | Module Size | Z-Summary Value |
| --- | --- | --- |
| HFD_015 | 50 | 0.7757 |
| HFD_016 | 41 | 0.9235 |
| HFD_011 | 67 | 0.9949 |
| HFD_012 | 63 | 1.3316 |
| HFD_000 | 655 | 2.4595 |
| HFD_014 | 53 | 3.5782 |
| HFD_005 | 117 | 5.0454 |
| HFD_010 | 76 | 5.1887 |
| HFD_013 | 54 | 5.3209 |
| HFD_017 | 40 | 8.8512 |
| HFD_007 | 85 | 10.95 |
| HFD_003 | 186 | 13.963 |
| HFD_009 | 82 | 14.075 |
| HFD_008 | 83 | 15.137 |
| HFD_001 | 323 | 16.222 |
| HFD_004 | 139 | 18.12 |
| HFD_006 | 97 | 18.121 |
| HFD_002 | 289 | 22.371 |

**B**
| HFD | C57WT | Adjusted P-Value |
| --- | --- | --- |
| HFD_002 | C57WT_002 | 5.19E-34 |
| HFD_000 | C57WT_000 | 2.14E-32 |
| HFD_007 | C57WT_013 | 1.76E-31 |
| HFD_003 | C57WT_008 | 9.87E-26 |
| HFD_006 | C57WT_006 | 6.87E-20 |
| HFD_001 | C57WT_005 | 5.72E-17 |
| HFD_004 | C57WT_003 | 7.25E-17 |
| HFD_008 | C57WT_017 | 4.51E-15 |
| HFD_009 | C57WT_016 | 4.74E-15 |
| HFD_011 | C57WT_012 | 1.67E-13 |
| HFD_005 | C57WT_020 | 3.71E-07 |
| HFD_010 | C57WT_014 | 1.25E-06 |
| HFD_013 | C57WT_007 | 0.000774 |
| HFD_016 | C57WT_023 | 0.007833 |

| HFD | C57MNX | Adjusted P-Value |
| --- | --- | --- |
| HFD_000 | C57MNX_000 | 6.38E-44 |
| HFD_008 | C57MNX_008 | 1.33E-40 |
| HFD_002 | C57MNX_001 | 2.19E-37 |
| HFD_012 | C57MNX_018 | 5.10E-33 |
| HFD_004 | C57MNX_003 | 3.38E-32 |
| HFD_001 | C57MNX_005 | 6.08E-22 |
| HFD_007 | C57MNX_007 | 2.25E-20 |
| HFD_003 | C57MNX_004 | 3.03E-18 |
| HFD_009 | C57MNX_002 | 8.79E-13 |
| HFD_010 | C57MNX_009 | 5.43E-09 |
| HFD_014 | C57MNX_015 | 3.92E-06 |

**Table S4:**
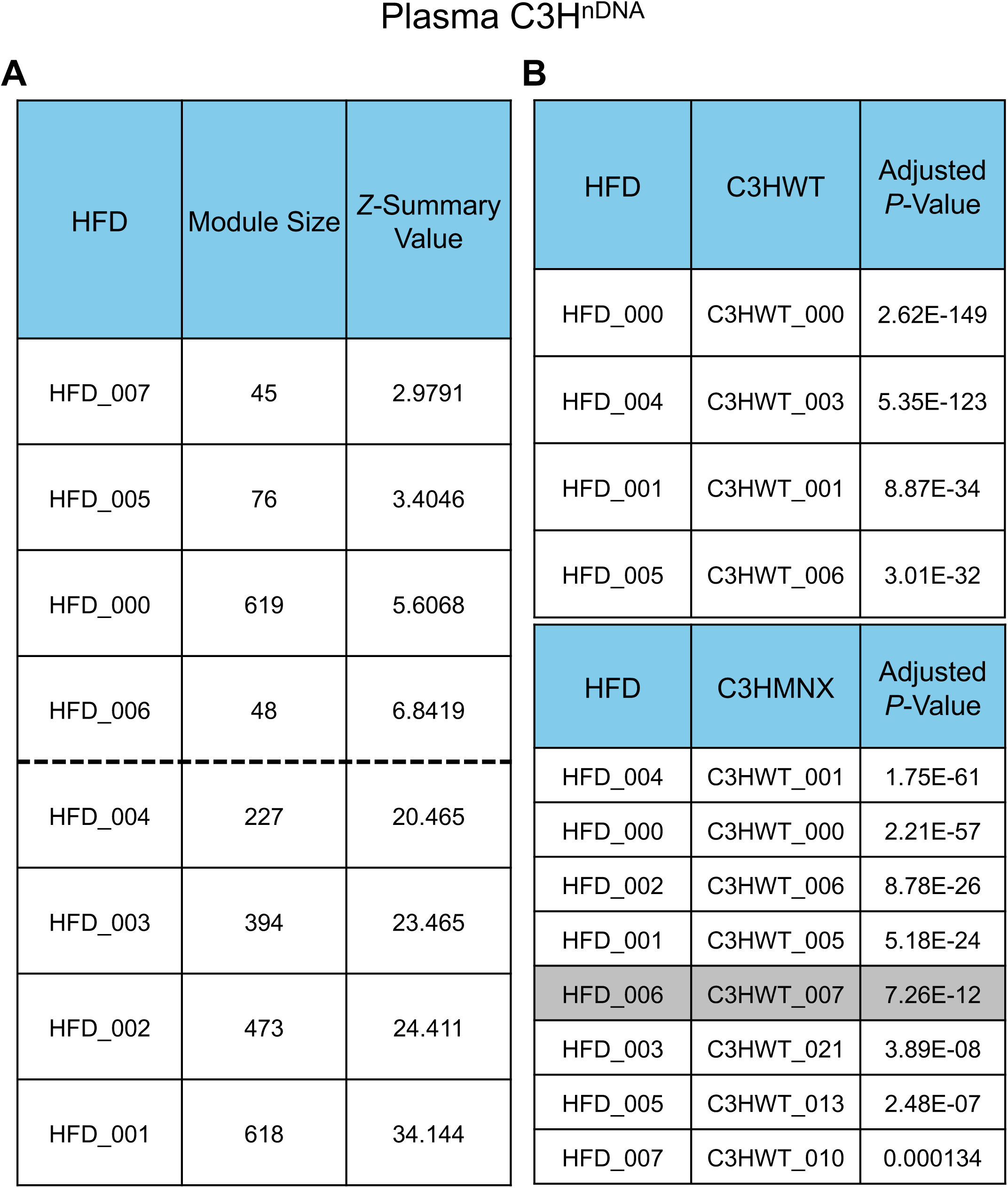
Results of overlap analysis and module preservation analysis within the Plasma HFD network in mice possessing C3H^nDNA^. (A) Results of module preservation analysis of High-Fat Diet (HFD) modules not found in Control Diet network. *Z*-Summary values are ranked and modules with values < 10 were retained. Module numbers are arbitrary labels. **(B)** HFD modules significantly associated with C3H^nDNA^:C3H^mtDNA^ WT (‘C3HWT’) and C3H^nDNA^:C57^mtDNA^ MNX (‘C3HMNX’) strains. Only HFD modules that are unique to the WT or MNX strain were investigated, not those that are shared by both strains. Adjusted *P*-value represents significance of association. Coloured modules are those represented in Figure 7.

